# Molecular Clock Dating of Deep-Time Evolution Using Complex Mixture Models

**DOI:** 10.1101/2025.07.17.665246

**Authors:** Sishuo Wang, Andrew Meade

## Abstract

Molecular clocks are a fundamental technique in evolutionary biology for establishing the timing and tempo of organismal divergence. However, currently available molecular clock methods, which often rely on simple homogeneous substitution models, can produce inaccurate time estimates, particularly for deep-time or rapidly evolving lineages where substitution heterogeneity and saturation are common. Hereby, we introduce phyloHessian (https://github.com/evolbeginner/phyloHessianWrapper), a Julia-based software to enable the use of complex mixture substitution models in molecular dating. phyloHessian computes the phylogenetic Hessian matrix and integrates it into PAML-MCMCtree’s approximate likelihood framework to conduct dating analyses. Simulations mimicking phylogenies at different timescales demonstrate that complex mixture substitution models significantly enhance the accuracy of divergence time and substitution rate estimates in deep-time phylogenies. This pattern remains consistent across a wide range of uncertainties associated with molecular clock analysis. Additionally, mixture models display greater robustness to model and calibration specifications compared to their homogeneous counterparts. Empirical analysis of ancient symbiont lineages Microsporidia and Rickettsiales with different substitution models shows that mixture models, compared to homogeneous models, yield accelerated substitution rates and in some cases significantly different divergence times, leading to a revised understanding of their host association origins. Our findings underscore the importance of incorporating complex mixture substitution models for constructing reliable evolutionary timelines and elucidating the evolutionary history of deep-time or fast-evolving lineages.

## Introduction

Historically, our understanding of evolutionary time scales relied on limited fossil records, until a significant change occurred in the 1960s when molecular clock theory was proposed (Zuckerkandl and Pauling 1965), making it possible to infer evolutionary time and rate using genetic data. While the original concept of the molecular clock assumed that the rate of gene sequence changes is constant among lineages, further evidence shows that even the same gene can exhibit variations in rate among different species, leading to the development of relaxed molecular clock models which allow for lineage-specific evolutionary rate and a series of derivative models (Thorne et al. 1998; Drummond et al. 2006; Yang and Rannala 2006). Over years, molecular clock theory has evolved, continuously increasing in complexity and accuracy, providing a powerful tool for studying biological evolution. Particularly, with the development of Bayesian phylogenetic methods and the rapid growth of genetic sequence data, Bayesian models based on Markov chain Monte Carlo (MCMC) methods have become the most widely used molecular clock dating method, which allows for a comprehensive examination of various uncertainties in analysis, including factors such as calibrations and across-lineage rate differences (Ho and Duchêne 2014; dos Reis et al. 2016).

While molecular clock approaches offer a powerful framework for inferring evolutionary timescales, the reliability of such inferences heavily depends on accurately modeling the evolution of the molecular data, such as protein sequences, alongside the incorporation of calibration information (Guindon 2020). Protein evolution is typically modelled as site-homogeneous process using empirically derived substitution matrices of amino acid exchangeabilities (e.g., LG or WAG) and fixed equilibrium frequencies, with variation in evolutionary rates modeled with a gamma distribution (Yang 1993). However, while computationally efficient, such models oversimplify real-world biological processes by failing to capture the heterogeneity in amino acid preferences driven by diverse structural and functional constraints across sites. To address these issues, site-heterogeneous substitution models have been proposed since (Lartillot and Philippe 2004; Pagel and Meade 2004). These models account for among-site substitution heterogeneity by incorporating predefined substitution matrices for different secondary structures and solvent accessibility (Le et al. 2008; Le and Gascuel 2010), across-site rate variation (Le et al. 2012), or mixtures of amino acid frequency profiles (Quang et al. 2008; Wang et al. 2008). Of particular interest is the last category, which captures different amino acid preferences across sites, as represented by the CAT model (Lartillot and Philippe 2004) and its simplified counterparts for maximum-likelihood phylogenetics called Cxx (where xx indicates 10 to 60 profile categories) (Quang et al. 2008). By better approximating the true biological complexity of amino acid substitution, profile mixture models mitigate artifacts like long branch attraction, thereby enhancing the ability to distinguish phylogenetic signal from homoplasy and displaying higher sensitivity to substitution saturation (Wang et al. 2018; Wang et al. 2019; Schrempf et al. 2020; Baños, Susko, et al. 2024). These features make profile mixture models especially well-suited for studying microbial evolution, where deep divergences and rapidly evolving sites are prevalent. Indeed, empirical analyses of deep-time phylogenomic data show that profile mixture models often exhibit superior model fit compared to homogeneous alternatives (Williams et al. 2020; Bujaki and Rodrigue 2022).

A particular outcome of using site-homogeneous models like LG+G is that branch lengths are usually underestimated in case of extensive sequence divergence (Venditti et al. 2008; Moody et al. 2022). This happens because proteins may not use all 20 amino acids at each position and homogenous models incorrectly interpret this restricted use of amino acids as evidence of less evolutionary change than what truly occurred over long time (Lartillot et al. 2007; Wang et al. 2008). To see this, consider a site restricted to Asp and Glu. A mixture model would identify a preference for negatively charged amino acids and infer unobserved substitutions (e.g., Asp → Glu → Asp) even if two sequences show the same amino acid, leading to longer branch lengths. Obviously, this pattern is more often observed with an increasing evolutionary scale. Studies have shown that the deepest branches of the tree of life, spanning approximately 4.0 billion years (Gyr), can display branch lengths up to twice as long under profile mixture models compared to the LG+G model (Moody et al. 2022; Wang and Luo 2025). Given that in molecular clock analysis, branch length, the expected substitutions per site, is the product of time and (average) substitution rate of that branch (dos Reis et al. 2016; Guindon 2020), an important question emerges: do profile mixture models, which often excel in deep-time phylogenetic inference, lead to more accurate divergence time and substitution rate estimates?

To our knowledge, none of the available molecular clock programs have directly implemented complex substitution models including those profile mixture models, with PhyloBayes’ CAT model being the exception—yet its high computational demands may limit its use in large-scale molecular clock analyses. We recently developed bs_inBV (Wang and Luo 2025), enabling the use of diverse substitution models in the popular MCMCtree molecular dating software by applying a bootstrap approximation of the Hessian matrix, which captures likelihood surface curvature and is required by MCMCtree’s fast approximate likelihood dating procedure (Thorne et al. 1998; dos Reis and Yang 2011). However, the bootstrapping process is computationally intensive, and its approximation sometimes inadequately represents the true Hessian. Additionally, and perhaps more importantly, the relative performance of different substitution models in molecular clock dating remains unexamined through systematic simulation studies, leaving a critical gap in our understanding of how choice in substitution model affects divergence time estimation, particularly in deep-time or fast-evolving phylogenies.

Hereby, we introduce phyloHessian, a Julia-based software that uses numerical methods to calculate accurate Hessian matrices for diverse complex substitution models and integrates them directly into MCMCtree’s approximate likelihood framework for rapid molecular clock dating. We perform extensive analyses on simulation data to comprehensively compare the performance of different substitution models in divergence time and substitution rate estimation under a wide range of conditions that mimic phylogenies from relatively recent to very ancient divergences. We further analyze empirical data to assess how different substitution models affect our understanding of ancient symbiont evolution and the origins of their host associations.

## Results

### phyloHessian: enabling molecular clock dating in MCMCtree with complex substitution models

To address the challenges in molecular clock analysis under various substitution models, we developed a Julia-based software phyloHessian. phyloHessian first calls IQ-Tree (Wong et al. 2025) or PhyML (Guindon et al. 2010), two widely used phylogenetic reconstruction software that implement a range of mixture models, to calculate the maximum-likelihood estimates (MLE) of branch length *θ̂* = (*θ̂*, …, *θ̂*)^T^ under user-specified substitution model. Let *θ* = (*θ*_1_, …, *θ*_*n*_) be the branch lengths and Δ*θ* = *θ* − *θ̂*. Next, it applies finite difference numerical methods to calculate the gradient 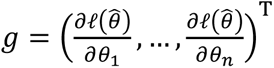 and Hessian matrix 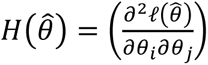 i.e., the first- and second-order derivatives of the log-likelihood of the phylogenetic data evaluated at the MLEs.. Last, the resulting Hessian matrix is integrated into MCMCtree to be used with its approximate likelihood method (Thorne et al. 1998; dos Reis and Yang 2011) for divergence time and rate estimation. MCMCTree’s approximate likelihood method approximates the log-likelihood surface for branch lengths using a Taylor expansion to the second order

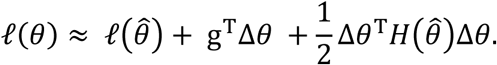

To demonstrate the reliability of phyloHessian, we compared its results with those obtained from well-established phylogenetics software packages. First, we found that phyloHessian and IQ-Tree produced identical log-likelihoods under all tested complex substitution models available in IQ-Tree (Data S1). Further, we validated phyloHessian’s Hessian calculation by confirming that the Hessian matrix obtained by phyloHessian and CODEML under site-homogeneous models (such as LG+G) were nearly identical (Data S2).

### Profile mixture substitution models enhance time and rate estimation in deep-time and fast-evolving phylogenies: evidence from simulations

To evaluate the impact of mixture substitution models (Materials and Methods: Mixture substitution model) on divergence time and substitution rate estimates, and to compare their performance to profile homogeneous models like LG+G, we conducted extensive simulations. To simulate a range of evolutionary scenarios spanning relatively recent to very deep-time divergences, we followed our prior study (Wang and Luo 2025) to generate 30 replicates of 20-tip birth-death trees with protein sequences that were 300 amino acids (aa) in length simulated under the substitution model LG+C60+G4{1.0} for both autocorrelated-rates (AR) and independent-rates (IR) molecular clock rate model. For simplicity, we use “+G” to represent “+G4{1.0}” hereafter [IQ-Tree’s default; four-category discrete Gamma distribution for the relative rate, shape parameter α = 1.0, i.e., Gamma(1,1)]; see also Materials and Methods: Simulation procedure. We mainly focused on five calibration strategies. These five calibration strategies consisted of a single calibration at the root with both upper and lower bounds (*root_only*), fully calibrated root plus one fully-bounded internal node (*single_interval*), fully calibrated root plus one minimum-bounded internal node (*single_min*), fully calibrated root plus two fully-bounded internal nodes (*two_interval*), and fully calibrated root plus two minimum-bounded internal nodes (*two_mins*).

Under the focal simulation scheme (see Materials and Methods: Simulation procedure), the profile mixture model LG+C60+G performed comparably to the traditional LG+G model for divergence time estimation when simulated root ages were set to either 1.0 or 2.0 billion years ago (Ga). However, for root ages ≥3.0 Ga, LG+C60+G obtained significantly more accurate time estimates than LG+G, with the performance gap becoming increasingly pronounced at 4.0 Ga (Fig. 1A). For example, under the *root_only* calibration strategy, LG+C60+G demonstrated superior time estimation accuracy (mean relative difference ± SD: 0.157 ± 0.061 for AR clock model, 0.139 ± 0.046 for IR clock model) compared to LG+G (AR: 0.193 ± 0.062; IR: 0.197 ± 0.061). Mean relative difference is defined as mean 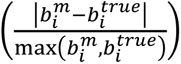, where 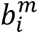 and 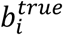 respectively denote the inferred posterior mean age and the true value for node *i*. The largest discrepancies in divergence time estimation occurred under the calibration scheme *two_min*, where LG+C60+G showed lower error than LG+G (mean relative difference: AR 0.160 vs. 0.232; IR 0.147 vs. 0.259; Fig. 1A). In contrast to divergence time estimates, LG+C60+G provided much more accurate branch-specific substitution rate estimates than LG+G across all tested root ages (Fig. 1B). The error reduction obtained by LG+C60+G compared with LG+G, calculated as 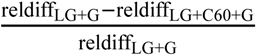 (i.e., the error reduction was calculated using the node-averaged mean relative differences between models), averaged across all calibration strategies, ranged from 26% to 53% under AR, and 23% to 55% under IR for root ages of 1.0 to 4.0 Ga (Table 1). The pattern held true with alternative metrics (mean branch score distance [BSD] or scatter-plot comparisons) for comparison with the simulation values (Figs. S1-S3).

**Figure 1.**
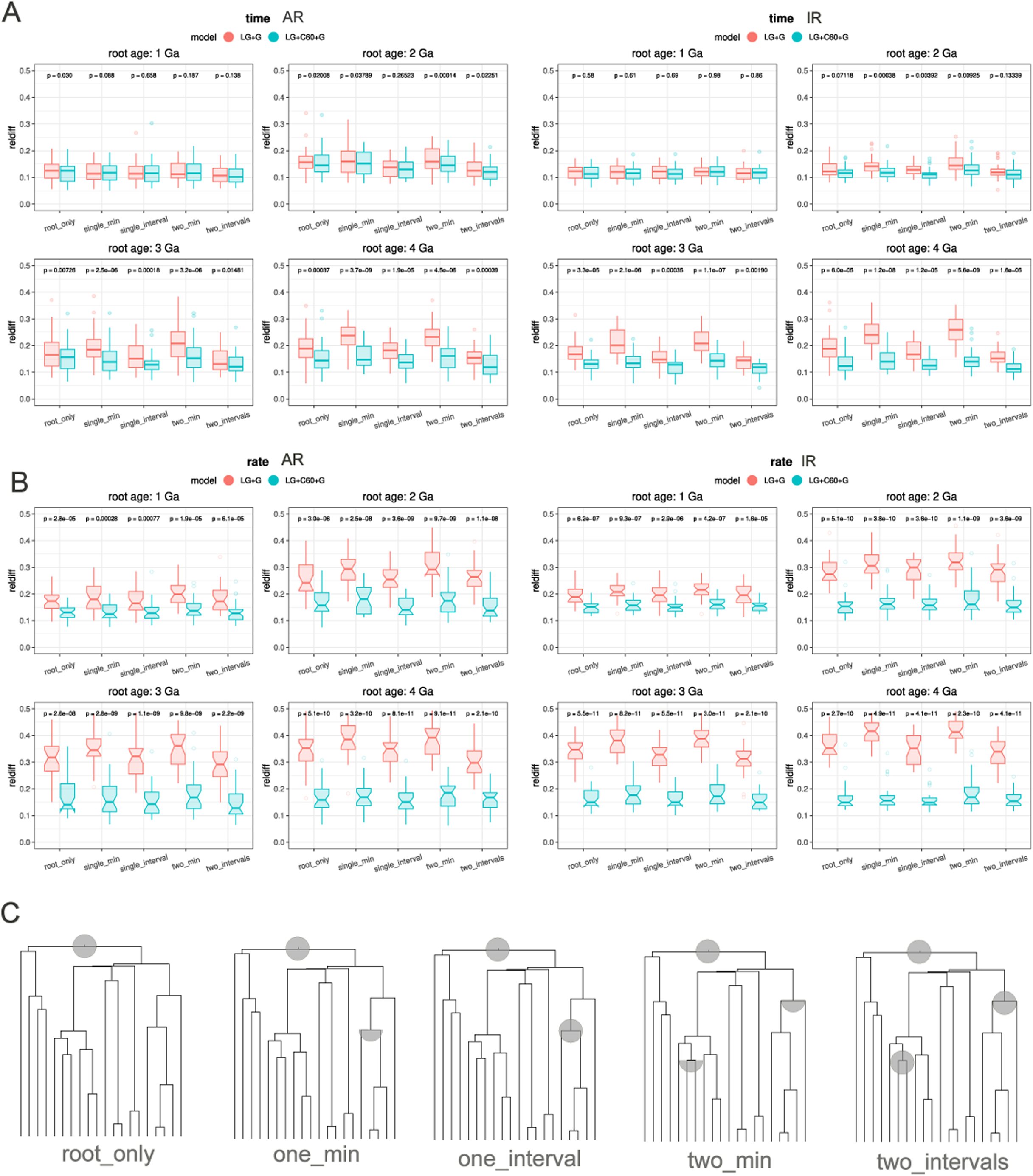
Comparison of the accuracy of divergence time and branch-specific substitution rate estimation under substitution models LG+G and LG+C60+G by simulation. (A, B) Each plot contains 30 dots representing the mean relative difference (compared to the true values used in simulation) based on MCMCtree molecular clock dating with the corresponding calibration strategy on 30 simulated datasets. For each simulation, 30 timetrees each containing 20 tips are simulated using a birth-death process with a mean substitution rate 0.25 substitutions/site/Gyr across four different root ages ranging from 1.0 to 4.0 Ga. Sequence alignments are generated with LG+C60+G4{1.0} (+G4{1.0} indicates α = 1 in a four-category discrete Gamma-distributed across-site relative rates) under AR (left-hand side) and IR (right-hand side) clock models respectively. *P*-values are obtained from a paired Wilcoxon signed-rank test. The box plots (median, interquartile range, 1.5×IQR whiskers) illustrate the relative differences in time (A) and rate (B) estimates compared to true values. (C) A schematic figure showing the five calibration schemes used for molecular clock analyses on simulated data. *root_only*: a single bounded calibration (both upper and lower bounds); *single_min*: bounded calibration at the root and one minimum-bounded internal node; *single_interval*: bounded calibration at both the root and an internal node; *two_min*: bounded calibration at the root and two minimum-bounded internal nodes; *two_intervals*: bounded calibration at both the root and two internal nodes.

**Table 1.**
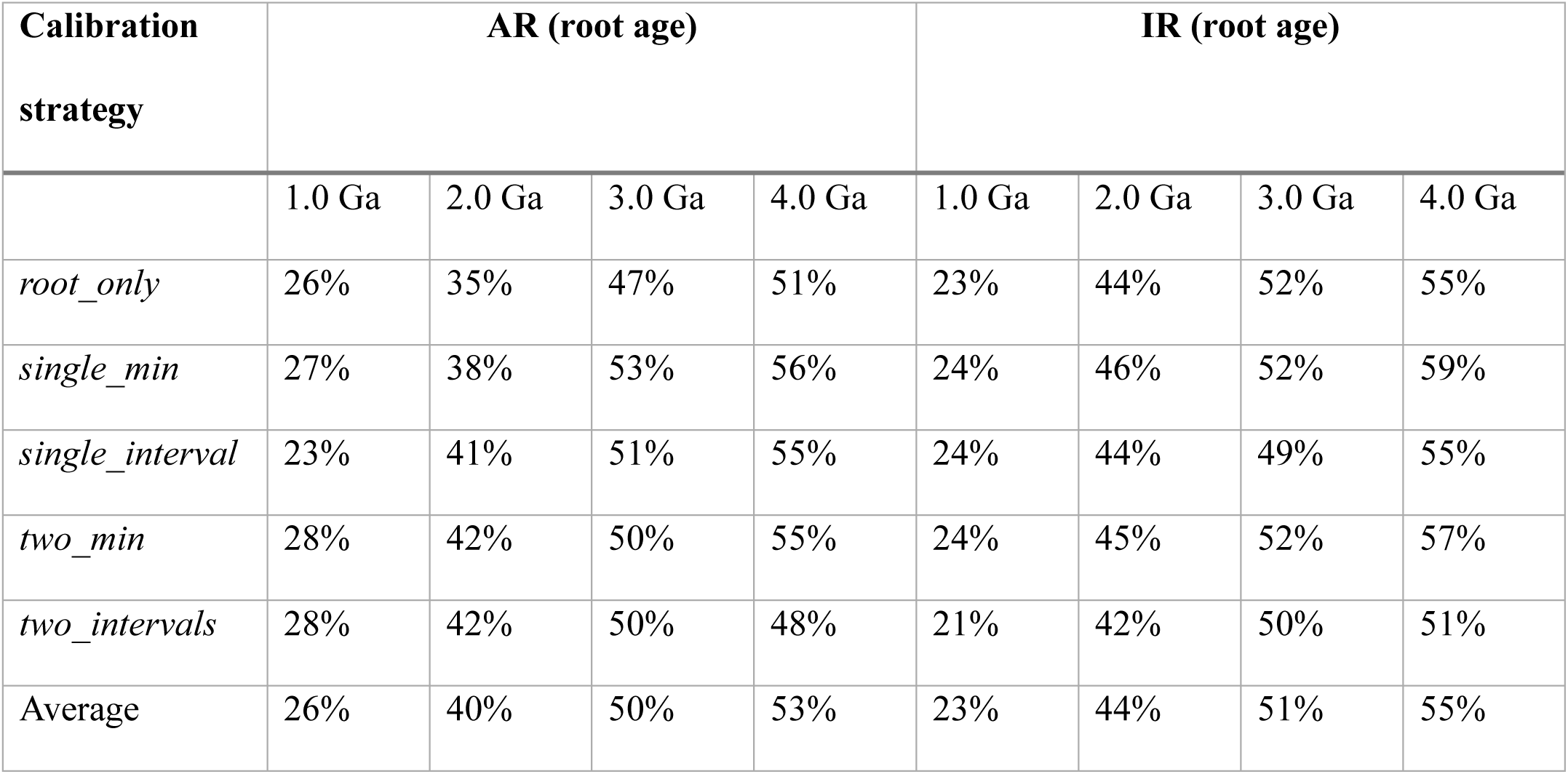
Reduction of the relative differences in substitution rate estimates, calculated as 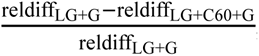, under LG+G versus LG+C60+G under different root ages and calibration strategies.

Next, we explored the patterns for more complex mixture models. We combined structurally constrained substitution models EX2 (exposed/buried sites) and EX3 (highly exposed/intermediate/buried sites) (Le et al. 2008) with the predefined equilibrium frequency C20 to account for site-specific variation in amino acid profiles by 20 categories of equilibrium frequencies. This resulted in 40 and 60 different rate matrices for EX2 and EX3 respectively. When simulation was done under profile-exchangeability mixture models EX2+C20+G and EX3+C20+G, the above patterns became more pronounced (Fig. S4). In contrast to profile mixture models, when alignments were simulated under exchangeability mixture models (EX2+G, EX3+G, and LG4M), LG+G and the true simulation model performed similarly in estimating divergence times (Fig. S5).

Large sequence divergence may arise from not only long divergence time, but also rapid evolution. To test whether different time estimates are affected by varying evolutionary rates (Fig. S6), we simulated alignments on phylogenies with a fixed root age of 1.0 Ga but different mean absolute substitution rates along the phylogeny (0.25-1.0 substitutions/site/Gyr). This represents evolutionary rates from relatively low, as in most microbes, to very fast, as observed in some pathogens. We would anticipate comparable sequence divergence whether we fixed the root age at 1.0 Ga and increased the average substitution rate (from 0.25 to 1.0 substitutions/site/Gyr; Fig. S6), or fixed the substitution rate at 0.25 substitutions/site/Gyr and increased the root age to 4.0 Ga (Fig. 1). Averaged across all calibration strategies, LG+C60+G displayed error reductions in substitution rate estimate of 65% (AR) and 61% (IR) at a mean substitution rate of 1.0 substitutions/site/Gyr.

### Performance of profile mixture models in molecular clock dating under varied simulation conditions

Next, we evaluated the influence of multiple sources of uncertainties in molecular clock dating on the performance of profile mixture and traditional homogenous models when sequence alignments are simulated under profile mixture models. These involve sequence length, Gamma-distributed across-site rate, birth-death process, variance in lineage-specific rates, and number of sequences (see Note S1 for details).

We first investigated how sequence length (ranging from 100 to 2000 aa) affected the accuracy of the estimates using different substitution models (Fig. S7). Although the increase in divergence time estimate accuracy achieved by using LG+C60+G instead of LG+G was minimal for 100-aa alignments, the improvement became increasingly apparent as the alignment length extended to 2000 aa. Across all analyzed calibration strategies, the LG+C60+G model consistently improved the accuracy of time estimation for the AR clock models compared to LG+G. For a simulated root age of 4.0 Ga, this improvement was substantial, resulting in an error reduction (see above) of 30%, 28%, 25%, and 12% for alignment lengths of 2000, 1000, 500, and 100 aa, respectively; the benefit was still significant but less pronounced for a younger root age of 3.0 Ga, with corresponding improvements of 22%, 18%, 15%, and 7%. Both LG+C60+G and LG+G models achieved higher estimation accuracy with longer sequences (i.e., lower values in *y*-axis from 100 to 2000 aa), indicating that longer sequences are more informative for more accurate molecular clock analyses, consistent with the notion that mixture models perform much better on longer alignments for phylogenetic reconstruction (Baños, Susko, et al. 2024). Regarding the substitution rate, mixture models displayed more accurate estimates across time scales for different sequence lengths (Fig. S8).

We further adjusted the gamma correction parameter (α) to modulate the variance of the across-site relative rates. Specifically, we set α to 0.5 and 1.5, which scaled the across-site variance to 2-fold (0.5/0.5²) (scheme *Gamma_0.5* in Fig. 2A) and 2/3-fold (1.5/1.5²) (scheme *Gamma_1.5* in Fig. S9A), respectively, compared to our focal analysis (α=1). These adjustments did not alter the patterns observed in the original analysis. Similarly, when exploring alternative birth-death process parameters, a flat birth-death process (simulation scheme: *flat_time_prior*) produced results comparable to those from the focal analysis (Fig. 2B). Under complete taxon sampling, the difference in time estimates between the LG+G and LG+C60+G models became more obvious (scheme *complete_taxon_sampling* in Fig. S9B). Moreover, increasing the standard deviation of the among-branch sequence absolute substitution rate to twice and three times its default value reduced the differences in divergence time estimation between using LG+G and LG+C60+G, despite the unchanged overall pattern (schemes *SD_lograte_0.38, SD_lograte_0.55* in Figs. 2C, S9C). In addition, although all simulations above were performed on 20-tip trees for simplicity, the analysis was extended to 40-, 60-, 80-, and 100-tip trees, with similar patterns found (Fig. S10).

**Figure 2.**
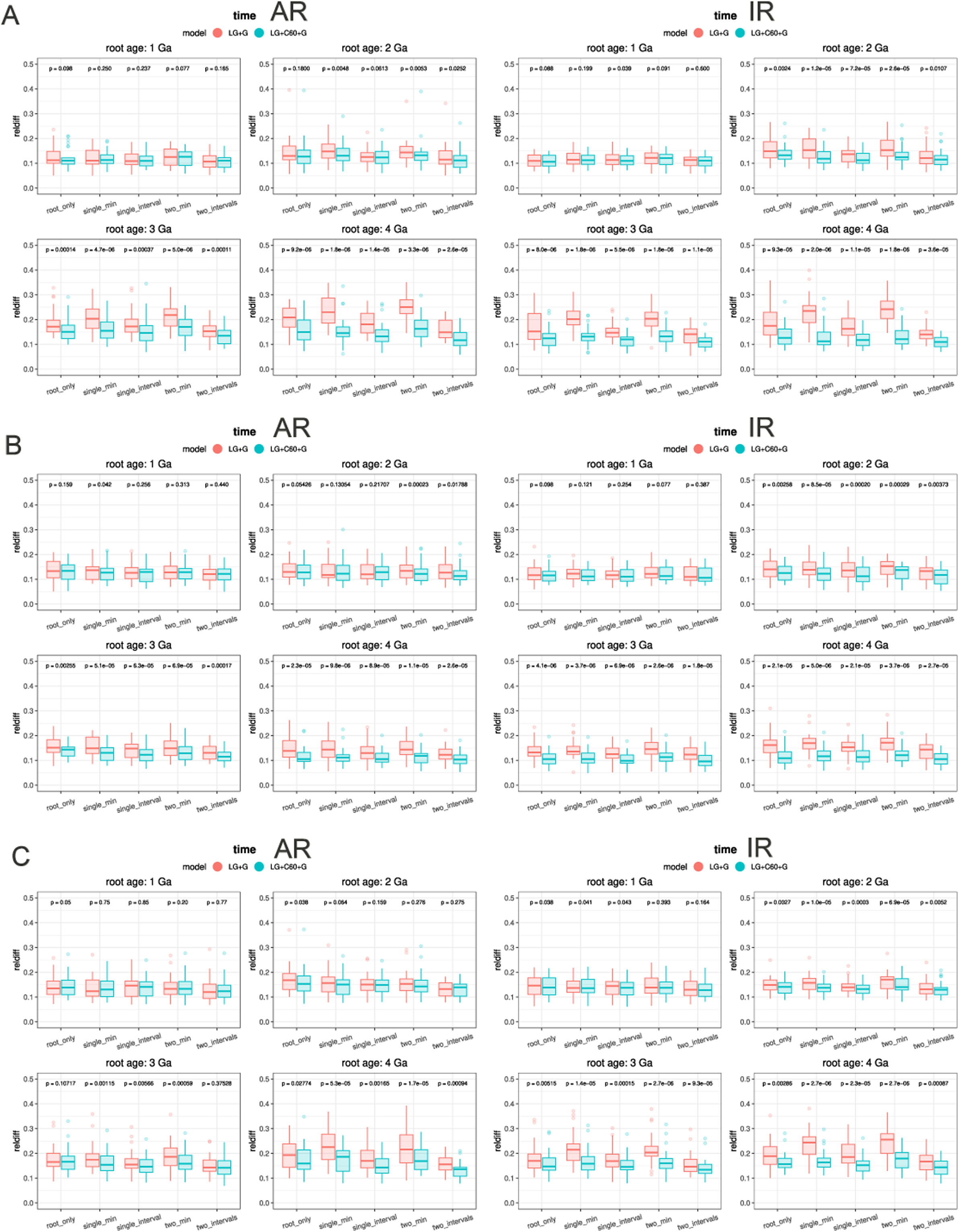
Investigating the robustness of divergence time estimation to alternative settings in simulation. (A) Sequence simulated under substitution model LG+C60+G4{0.5} (*Gamma_0.5*; the parameter alpha in the discrete Gamma distribution equal to 0.5), thus more rate heterogeneity across sites than the focal analysis (LG+C60+G4{1.0}). (B) Timetree simulated under a flat time prior (*flat_time_prior*). (C) Timetree simulated under a molecular clock with larger across-branch rate variance (*var_lograte_0.38*). The plots on the left and right correspond to analyses using the AR and IR clock models, respectively. More detailed information about the above alternative simulation schemes can be found in Note S1. Calibration strategies and other settings are the same as those used in Fig. 1.

### Profile mixture models display higher robustness to model misspecification

Further, we applied simulations to test the robustness of profile mixture models to misspecifications in calibrations or substitution models. To explore whether mis-specified calibrations would affect estimation under different substitution models when the sequences were simulated under LG+C60+G, we calibrated the nodes by adjusting their ages. These adjustments consisted of fixed percentage increases (shifting ages toward the past) or decreases (shifting ages toward the present) (Fig. 3). In general, LG+C60+G achieved higher accuracy when calibrations were older than the true divergence times (Fig. 3B). For a root age of 4.0 Ga and across all tested calibration strategies, on average, the relative difference in time estimation compared to true values was: 0.256 (LG+G) versus 0.195 (LG+C60+G) for the AR model, and 0.278 (LG+G) versus 0.185 (LG+C60+G) for the IR model. For root age of 3.0 Ga, these values changed to 0.252 (LG) versus 0.202 (LG+C60+G) for AR, and 0.255 (LG) versus 0.192 (LG+C60+G) for IR (Fig. 3B). Subtle differences between using LG+G and LG+C60+G were observed for root age ≤ 2.0 Ga. However, when calibrations were biased towards younger ages, the LG+G and LG+C60+G models generally yielded comparable accuracy for divergence time and substitution rate estimates, irrespective of the root age, with significant differences arising occasionally.

**Figure 3.**
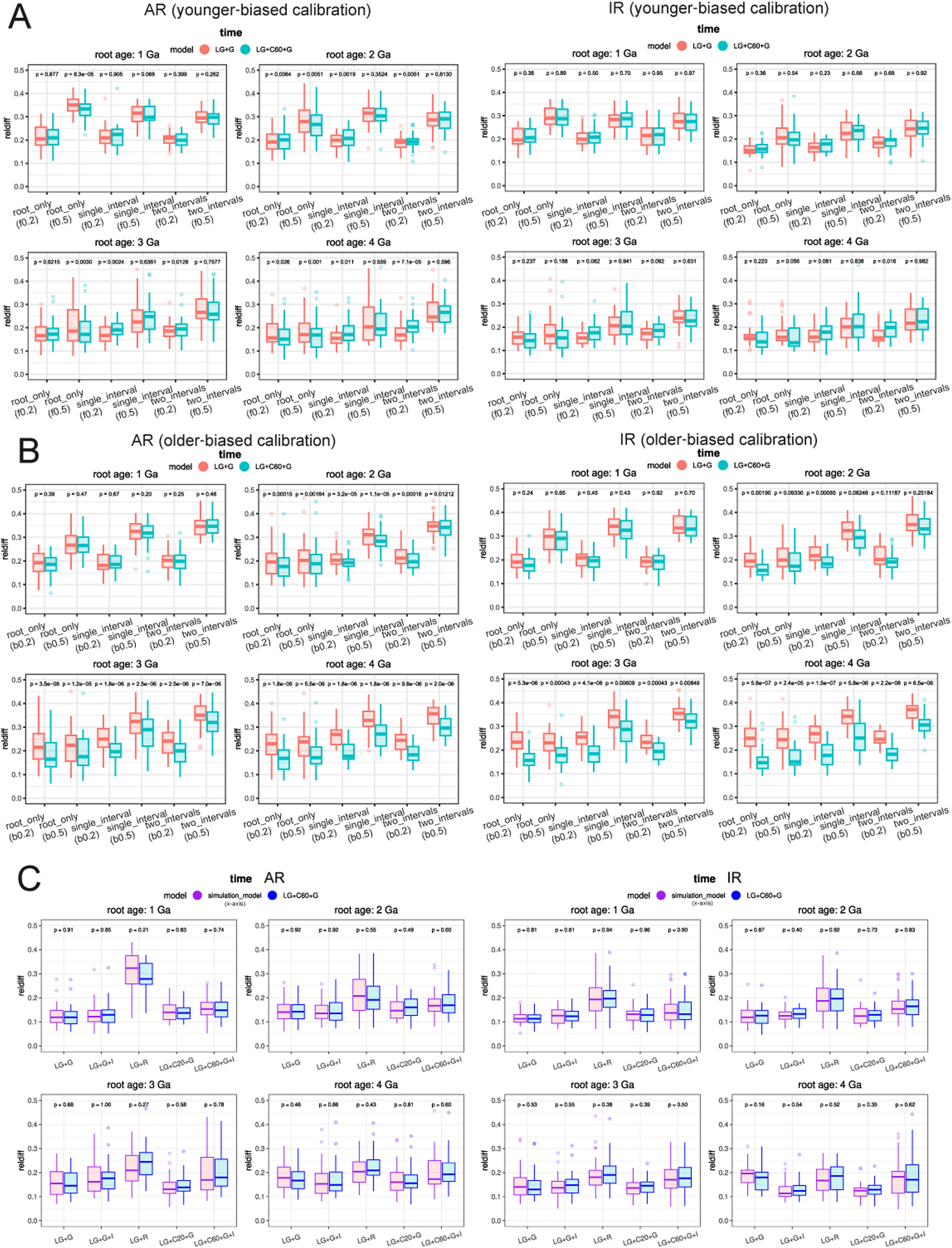
Evaluation of the impacts of calibration and model misspecification on molecular clock dating. Accuracy of divergence time estimation (relative difference) is shown as the *y*-axis. (A, B) The lower and upper time bounds of all calibrations are shifted forward (A) and backward (B) in time by 20% (f0.2, b0.2) or 50% (f0.5, b0.5) of their true values, respectively. Three calibration schemes are considered: (i) a root-only calibration (*root_only*), (ii) the root plus one fully calibrated internal node (*single_interval*), and (iii) the root plus two fully calibrated internal nodes (*two_intervals*), each with time ranges of the true value ±20% or ±50%. (C) Assessment of the accuracy of divergence time estimation under alternative substitution models used in simulation (LG+C60+G used in most other analyses): LG+G, LG+G+I, LG+R, LG+C20+G, LG+C60+G+I. Boxplots compare the accuracy obtained by the true model (purple) and by LG+C60+G (blue). Each plot contains 30 dots representing the mean relative difference based on MCMCtree molecular clock dating with the corresponding calibration strategy on 30 simulated datasets. For each simulation, 30 timetrees each containing 20 tips are simulated using a birth-death process with a mean substitution rate 0.25 substitutions/site/Gyr across four different root ages ranging from 1.0 to 4.0 Ga. The plots on the left and right correspond to analyses using the AR and IR clock models, respectively. Calibration strategies are the same as those used in Fig. 1.

The above analyses focused on alignments simulated under complex substitution models, particularly LG+C60+G. To determine how these models perform in molecular clock analysis when sequences evolve under simpler substitution models, we performed additional simulations using these models and the previously described settings (varying the simulated root ages from 1.0 to 4.0 Ga (Fig. 3C). These analyses indicate that the profile mixture model LG+C60+G displayed robust time and rate estimation, regardless of the true underlying substitution process. The robustness of LG+C60+G is evident in the following comparisons. When sequences were simulated under the basic LG+G model, LG+C60+G provided estimates of divergence times and absolute substitution rates at least as accurate as those obtained using LG+G. Further, similar results were observed when simulations were conducted with other models with factors not explicitly modeled by LG+C60+G, such as invariant sites (+I) and free rate across-site heterogeneity (+R) (Note S1). These findings suggest that LG+C60+G’s robustness was maintained even when the true evolutionary process involves additional complexities in substitution models.

### Comparison between profile mixture models with PMSF (posterior mean site frequency) and bootstrapped-Hessian approximations in molecular clock dating

The computational intensity of the LG+C60+G model often necessitates the PMSF approximation to accelerate phylogenetic reconstruction, especially for large datasets (Wang et al. 2018). This occurs because PMSF requires the computationally intensive pruning algorithm to run only once per site pattern using an approximated equilibrium frequency for each site, versus *c* times for a mixture model with *c* classes. A distinct approximation, previously employed in molecular clock analyses under complex substitution models, uses a bootstrap-based method to approximate the Hessian matrix and is implemented in the software bs_inBV (Wang and Luo 2025). To assess the impact of these two approximations on molecular clock dating under profile mixture models, we compared both divergence time and substitution rate estimates obtained under these approximations by evaluating their relative differences from the true simulation parameters.. We found that both approximations provided accurate estimates of divergence times, no matter which root age was used in simulation (Fig. 4A). In terms of the time estimates, across all simulated root ages, compared to LG+C60+G estimated rates, the average differences for AR were 3.6 ± 1.0% (PMSF approximation; *LG+C60+G+PMSF*), 7.1 ± 0.6% (bootstrapped-hessian; *LG+C60+G+bs_inBV*), and 7.3 ± 0.8% (combined use; *LG+C60+G+PMSF+bs_inBV*); for IR, the differences were 1.8 ± 0.4%, 5.9 ± 0.4%, and 6.5 ± 0.5%, respectively. The deviations in time estimates from these approximations, particularly PMSF, were close enough to the stochastic variation observed across independent MCMC runs of the original profile mixture model (*LG+C60+G_run2* in Fig. 4; comparisons in Fig. 4 are made relative to the initial MCMC run under LG+C60+G). The estimates of evolutionary rates exhibited slightly greater divergence from the original model with either approximation (Fig. 4B). The mean relative differences under both AR and IR models were around 5-10% for either approximation, or their combination, compared to the true parameters used in simulation.

**Figure 4.**
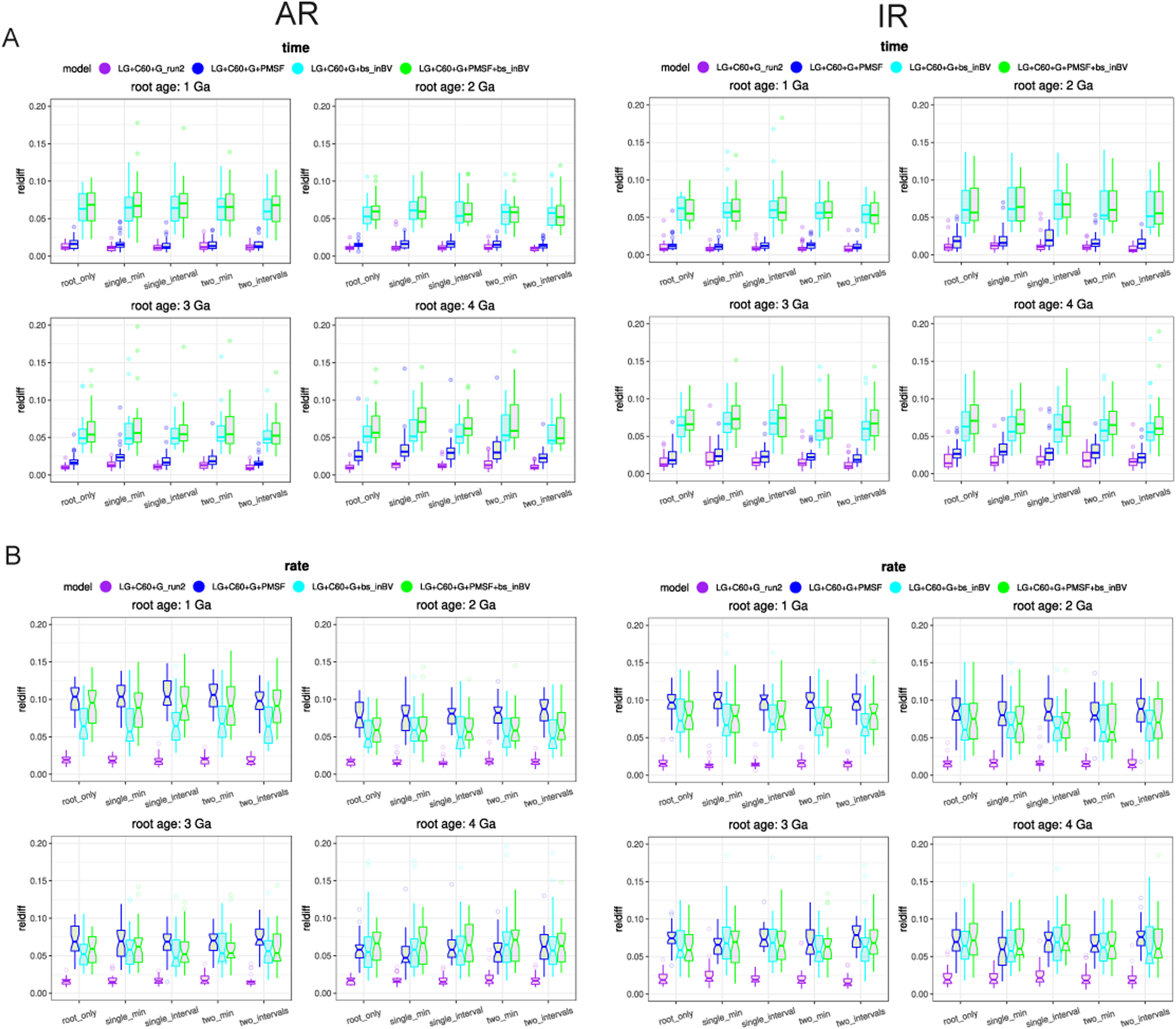
Evaluation of the approximation of PMSF and bootstrapped Hessian on molecular clock analysis by simulation. Phylogenies are simulated under the same strategy as the focal dataset. The sequence alignment is simulated under LG+C60+G{1.0}. The bootstrapped Hessian matrix is calculated using the software bs_inBV. The *y*-axis indicates the relative accuracy of the posterior mean of the divergence times (A) or substitution rates (B) compared to that obtained by the first MCMC run under LG+C60+G (note that this is different from other figures where the baseline in comparison is the true simulated times and rates). Each plot contains 30 dots representing the mean relative difference based on MCMCtree molecular clock dating with the corresponding calibration strategy on 30 simulated datasets. For each simulation, 30 timetrees each containing 20 tips are simulated using a birth-death process with a mean substitution rate 0.25 substitutions/site/Gyr across four different root ages ranging from 1.0 to 4.0 Ga. *LG+C60+G_run2*: the second independent MCMC run of MCMCtree under LG+C60+G. *LG+C60+G+PMSF*: PMSF approximation with LG+C60+G. *LG+C60+G+bs_inBV*: bootstrapped Hessian under LG+C60+G. *LG+C60+G+PMSF+bs_inBV:* bootstrapped Hessian under LG+C60+G+PMSF. Calibration strategies are the same as those used in Fig. 1.

We further compared the log-likelihood calculated by phyloHessian using Hessian computed by the finite-difference method (see Materials and Methods) and approximated via bootstrap with bs_inBV. Roughly speaking, both methods approximated the true log-likelihood of the data reasonably well. phyloHessian achieved higher Pearson correlation coefficients with the exact likelihood than the bootstrap-based method (*P*-value = 0.00026, paired Wilcoxon test; Fig. S11), on the data tested, while running ∼10 times faster (assuming 1000 bootstraps for bs_inBV). More details are given in Discussion.

### Divergence times and substitution rates under mixture models for the ancient pathogens Microsporidia and Rickettsiales: an empirical study

To evaluate mixture models on empirical data, we selected two ancient symbiont lineages with distinct evolutionary significance and accelerated evolutionary rates: Microsporidia, highly reduced, fast-evolving obligate intracellular parasites of animals related to fungi (Capella-Gutiérrez et al. 2012), and Rickettsiales, an alphaproteobacterial lineage that shares a close evolutionary relationship with the ancestors of mitochondria (Martijn et al. 2018; Wang and Luo 2021). Their rapid evolution combined with ancient divergence makes them ideal test cases for our analysis. Note that our primary goal was to assess how complex substitution models influence the molecular clock analysis itself, leading us to test different settings rather than selecting a single optimal configuration (e.g., partitions, clock model).

For both datasets, we partitioned the alignments into three subsets (see Materials and Methods) according to a Gaussian mixture model-based clustering according to the logarithm of the estimated substitution rates of each gene, and selected the best-fitting substitution models for each partition under both site-homogeneous and mixture substitution models respectively. In all cases, profile mixture models were associated with the lowest AIC values, with LG+Cxx+F+I+R4 (where xx = 40, 50, 60) being the most frequently selected (Data S3-S4).

Microsporidia do not have fossil records. Thus, we applied different calibrations to only the root node based on previous estimate of the occurrence time of its LCA (last common ancestor) (Note S2). We found that the time estimates for Microsporidia lineages were largely consistent between the best-fitting profile mixture and homogenous models (Fig. 5). The LCA of Microsporidia was estimated to occur at approximately 0.41 Ga (95% HPD: 0.74-0.45 Ga) and 0.63 Ga (95% HPD: 0.73-0.52 Ga) under the AR and IR clock models, respectively (Fig. 5A). In contrast to the similar divergence time estimates between homogeneous and mixture models, the absolute substitution rate was at least 1/3 higher under the mixture model (Fig. 5B). All microsporidian lineages experienced accelerated sequence changes compared to the outgroup lineages. While the highest substitution rate was predicted for the branch leading to the LCA of Microsporidia under the IR clock model, the AR clock model instead predicted the highest rate on the branch leading to *Enterocytozoon*, a major microsporidian pathogen in human and animals. The above pattern was consistent with alternative calibrations and different numbers of partitions (Fig. S12).

**Figure 5.**
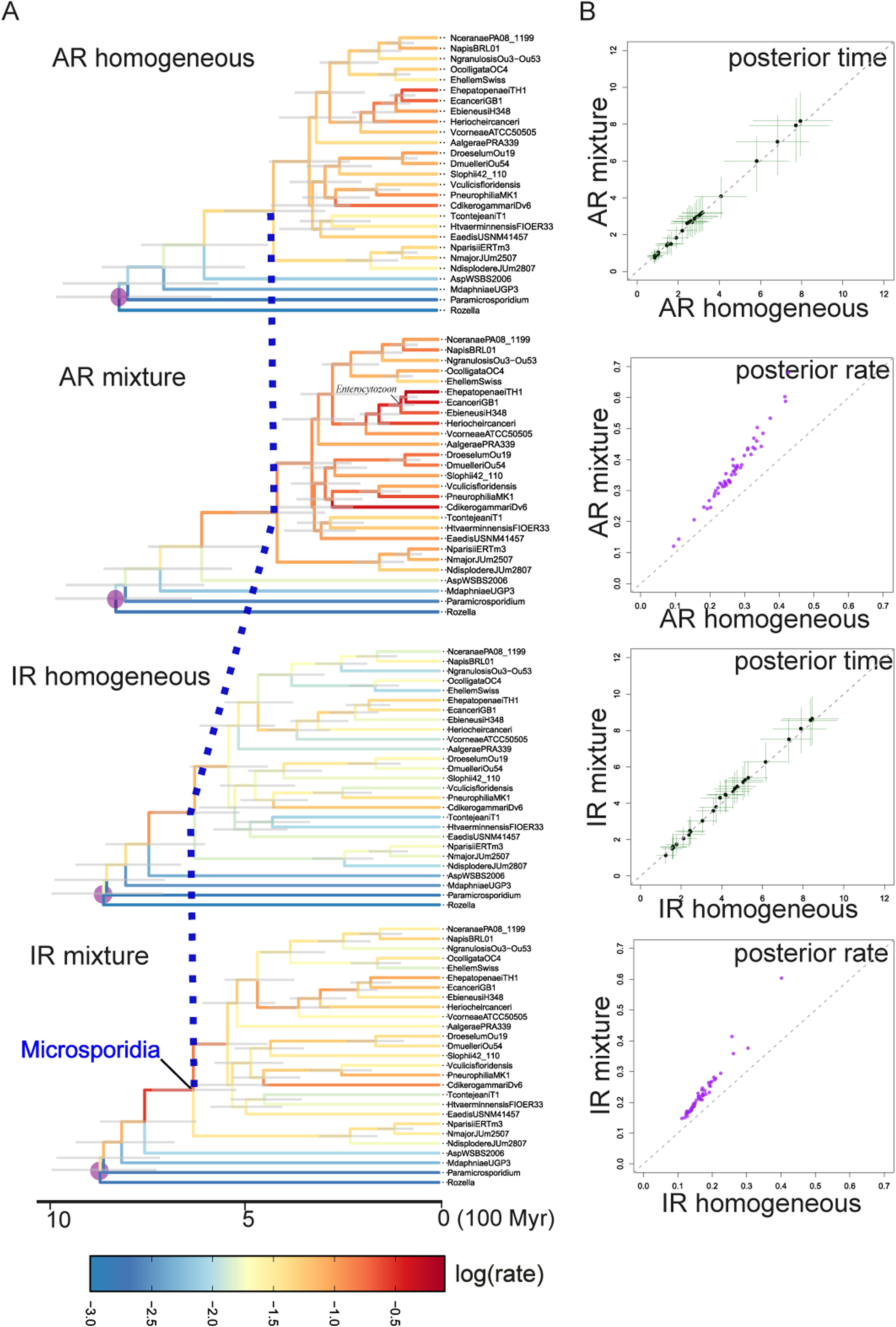
Evolutionary timeline and substitution rates estimated by MCMCtree under different substitution models for Microsporidia. A uniform calibration prior between 0.5 and 0.9 Ga was applied to the root (see Note S2), with soft bounds extending 2.5% beyond each time bound. Branch colors indicate the logarithm of the posterior mean substitution rate (estimated by MCMCtree), ranging from blue (low) to red (high). (B) Posterior divergence times and branch-specific substitution rates estimated by the mixture substitution model (*y*-axis) versus those estimated by the homogeneous substitution model (*x*-axis) under the AR or IR clock models. The bars indicate the 95% HPD.

Rickettsiales also lack direct fossil evidence; however, due to their close phylogenetic relationship with mitochondria, we employed the mitochondrial endosymbiosis dating strategy as used in our prior study (Wang and Luo 2021). This approach allows estimating Rickettsiales’ evolutionary timeline by incorporating eukaryotic fossils, based on the evolutionary scenario that mitochondria evolved from a lineage closely related to Alphaproteobacteria (and thus Rickettsiales). Particularly, under the AR clock model, the time estimates of Rickettsiales lineages under the mixture substitution model differed by an average of >20% compared to those from the homogeneous alternatives (Fig. 6A). In general, the ages of most lineages were estimated to be older under the profile mixture model (Fig. 6A). For example, under the AR clock model, the LCA of Rickettsiales was estimated to occur at 2.67 Ga (95% HPD: 2.97-2.35 Ga) by the mixture model, whereas the homogeneous substitution model estimated it at 2.26 Ga (95% HPD: 2.57-1.94 Ga). Similarly, eukaryotes were estimated to originate at 2.29 Ga (95% HPD: 2.57-1.98 Ga) under the mixture model, compared to 1.74 Ga (95% HPD: 1.98-1.50 Ga) under the homogeneous model (Fig. 6A). The above pattern holds under alternative calibration settings (Fig. S13). With a single partition (Fig. S13A, S13B), the difference in time estimates between different substitution models was less obvious. In all calibration schemes, the substitution rates under the mixture models were roughly 1.5 to 2 times as fast as those estimated under homogeneous models (Fig. 6B).

**Figure 6.**
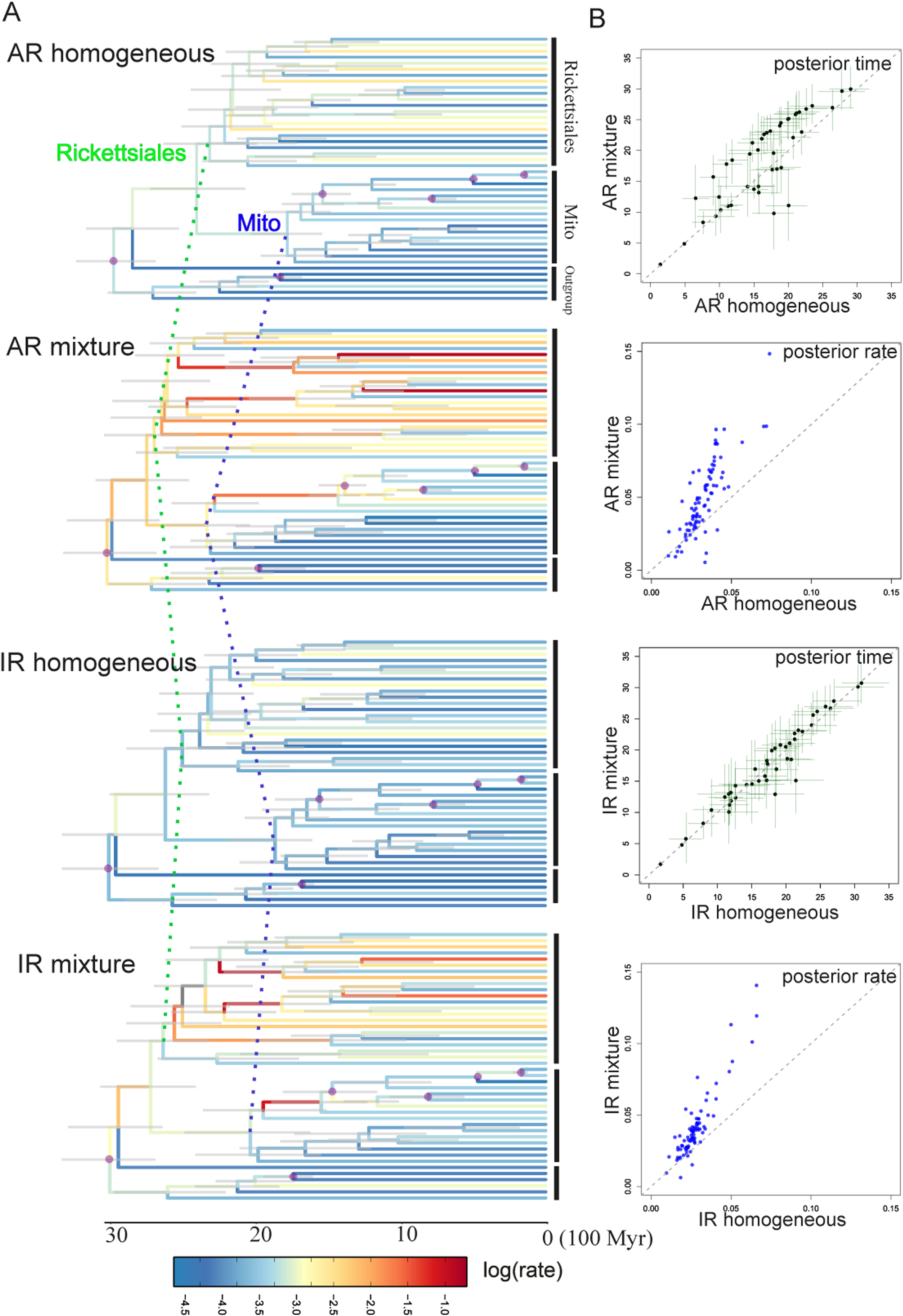
Evolutionary timeline and substitution rates estimated by MCMCtree under different substitution models for Rickettsiales. This is based the mitochondrial endosymbiosis dating strategy (Wang and Luo 2021), using eukaryotic fossils to calibrate Rickettsiales evolution, based on the evolutionary scenario that mitochondria originated from a lineage closely related to α-Proteobacteria. (A) The estimated evolutionary timeline of Rickettsiales under different substitution (the best-fitting homogeneous or mixture) and clock (IR or AR) models. All calibrated nodes are denoted by a circle. In addition to the calibration at the root, all four calibrations are placed within eukaryotes in the clade of mitochondria (see Note S2). Branch colors indicate the logarithm of the posterior mean substitution rate estimated by MCMCtree, ranging from blue (low) to red (high). (B) Posterior divergence times and branch-specific substitution rates estimated by the mixture substitution model (*y*-axis) versus those estimated by the homogeneous substitution model (*x*-axis) under the AR or IR clock models. The bars indicate the 95% HPD.

A recent study (Baños, Susko, et al. 2024) suggests that the empirical frequency estimated from the alignment (+F) might mislead phylogenetic reconstruction for profile mixture models. As an additional analysis, we re-performed molecular clock dating with the best-fitting mixture model without the empirical frequency option. Similar patterns were found (Figs. S14-S15).

## Discussion

### Approximate likelihood dating with complex models

The present study introduces phyloHessian, enabling Hessian calculation under diverse mixture and non-mixture amino acid substitution models for molecular clock dating using MCMCtree’s approximate likelihood method (Thorne et al. 1998; dos Reis and Yang 2011). A key question is whether MCMCtree’s approximate likelihood method performs equally well as exact likelihood under mixture substitution models in terms of divergence time estimation. While previous research suggests that this holds true for homogeneous models (dos Reis and Yang 2011), a full comparison is impossible due to the lack of software directly implementing complex substitution models. Therefore, we compared the phylogenetic likelihoods of bootstrap trees calculated by the approximate and exact likelihood methods under LG+C60+G using phyloHessian. The approximate likelihood values obtained from phyloHessian (using finite differences) generally showed good agreement with those derived from both the exact method and bs_inBV, despite minor discrepancies (Fig. S11). Further, phyloHessian sometimes outperformed bs_inBV, with bs_inBV showing markedly lower log-likelihood in occasional cases (Fig. S11). Discrepancies in time and rate estimates between the Hessian-based method (phyloHessian) and the bootstrap approximation (bs_inBV) stem from three factors (Fig. 4): the inherent limitation of bs_inBV due to the finite number of bootstrap replicates; the statistical assumption in bs_inBV that the gradient is zero when approximating the Hessian via the covariance matrix, an assumption that is invalid when the MLEs of branch lengths are near zero (dos Reis and Yang 2011); and the difference in parameter handling, as MCMCtree fixes substitution model parameters (such as the parameter α for the gamma distribution for relative rates) at their MLEs while bs_inBV re-estimates all parameters in each bootstrap, resulting in phyloHessian using only the branch length Hessian versus bs_inBV approximating the full parameter Hessian. A more thorough examination in time estimates, instead of the phylogenetic likelihood, is necessary to fully address the question in future.

It was when we were finishing writing this manuscript that we became aware, through personal communication, of another computational software, IQ2MC (Demotte et al. 2025), implemented in the newest version of IQ-Tree (v3.0.1). IQ2MC leverages the highly optimized computation of IQ-Tree’s phylogenetic library enabling efficient Hessian matrix estimation under not only amino acid, but nucleic acid and partitioned models, unlike phyloHessian which is restricted to amino acid models. Due to these computational advantages, IQ2MC was markedly faster than CODEML and phyloHessian (Data S5). However, our software, phyloHessian, offers more flexible settings in molecular dating by alternative Hessian calculation algorithms and interactions with other software like PhyML and bs_inBV. Its code for phylogenetic likelihood calculation under complex substitution models would be a valuable resource for phylogenetic software development within the Julia ecosystem.

Other methods, such as McmcDate (Mahendrarajah et al. 2023), also accelerate molecular dating analyses by approximating the phylogenetic likelihood. It is in concept similar to bs_inBV (Wang and Luo 2025), but operates in a Bayesian framework. A key advantage of bs_inBV and McmcDate over phyloHessian and IQ2MC is their ability to accommodate substitution models not yet available in specific phylogenetics software, such as FunDi (Gaston et al. 2011) or CAT-PMSF (Szánthó et al. 2023). However, these approximations via bootstrap or MCMC sampling may sacrifice computational speed and accuracy to some extent (Fig. 4), as discussed above.

### Deep-time molecular clock dating: the influence of complex mixture models

Our extensive simulations demonstrate that the advantage of using the complex mixture substitution model for divergence time estimation becomes increasingly significant as evolutionary timescales deepen (Figs. 1-4). Particularly, when the root age was earlier than 2.0 Ga, the difference became obvious. This highlights the importance of substitution models that better fit the data for dating deep-time phylogenies. On the other side, the substitution rate was almost always estimated higher and more accurate under mixture models than under homogeneous models. The underlying logic is straightforward. Mixture models typically estimate longer branch lengths than homogeneous models (Venditti et al. 2008; Moody et al. 2022; Wang and Luo 2025). Because branch length equals substitution rate multiplied by time (dos Reis et al. 2016; Guindon 2020), the underestimated branch lengths under simpler substitution models must be attributable to changes in either the rate, the time, or both. Given that molecular clock analyses typically constrain the time estimates through fossil-based calibrations, the increased branch lengths under mixture models primarily result in higher rate estimates, with a smaller impact on the time estimates themselves.

The case in empirical data analysis is more complex. Unlike simulations with known parameters, empirical data could arise from processes so complex that even the best-fitting models are inadequate. This notion is supported by the frequent selection of the most highly parameterized models as the best fit, implying that the complexity of the real process the sequence is generated far exceeds the capabilities of our current models. Consequently, they could make little difference in the time estimates compared to homogeneous models, because both might be an oversimplification of the true model. In such cases, it would be worth applying models like UDM (Schrempf et al. 2020) with more profile mixtures than C60, or the GTRpmix exchangeability matrix and its derived models (Baños, Wong, et al. 2024). While some of these models have been integrated into phyloHessian, a comprehensive evaluation of their applications in molecular dating analysis on empirical data remains a task for future research.

While the “mean relative difference” metric used to compare between models takes the absolute value, it is possible to tell a presumable bias in the time estimates from Figs. S2-S3: the posterior ages estimated by the homogeneous model seemed to overestimate the divergence times in scenarios mimicking deep-dating evolution in simulation. As such, it would be very important to have appropriate maximum time bounds on internal nodes (Wang and Luo 2021; Moody et al. 2022). If indeed in current molecular clock models there is a systematic bias in overestimating the divergence times, it might also explain the observation that mixture models displayed greater improvements in time estimation for alignment simulated under mixture models in the absence of maximum time bounds (*single_min*, *two_min*) compared to the case where internal nodes are fully calibrated (*single_interval*, *two_intervals*; Fig. 1A). In contrast to simulations, the Rickettsiales data reveals a contrasting pattern where mixture models led to older age estimates (Fig. 6). We suspect that one important factor lies in the calibration settings. In simulations, we have complete control over the calibration settings. Real-world data introduces complexities absent in simulation because the available calibrations are not uniform in their quality. While some might accurately reflect the true age of the node, others could be overly young, overly old, or simply uninformative. This complex interplay between substitution models and the quality and number of time calibrations, along with other variables such as the clock model and sequence length, likely makes the impact of mixture substitution models on posterior age estimates case dependent.

In summary, while the present study indicates significant impacts of complex profile mixture substitution models on divergence time inferences for deep-time or fast-evolving lineages, the importance of these models on time estimates cannot be overstated; their effect likely requires consideration on a dataset-specific basis. Therefore, we recommend rigorous molecular clock analyses employing different substitution models to assess their impact on divergence time and substitution rate estimates in future research. Note also that while our simulations used an LG+C60+G model, the LG exchangeabilities themselves were not estimated in the presence of profile mixtures (Le and Gascuel 2008). Ideally, for practical applications, it would be more appropriate to use models that directly estimate exchangeabilities under a corresponding profile mixture model for such cases (Baños, Wong, et al. 2024).

### Implications for the evolution of ancient symbionts

For empirical datasets of Microsporidia and Rickettsiales, profile mixture models, which display much higher model fit to data than their homogeneous counterparts, produced significantly higher substitution rates than estimated by homogeneous models (Figs. 5B, 6B). The difference suggests that bacterial pathogens/symbionts could experience relatively high evolutionary rates up to 1.0 or 2.0 Gyr. This hints that factors like elevated mutation rates, selection, or persistently reduced effective population sizes, which are often responsible for rapid sequence divergence, can have more profound and long-lasting effects on the evolutionary trajectory of symbiotic microorganisms than previously thought. Studying the substitution rates will also improve our understanding of microbial lifestyle evolution.

One of the merits of the mitochondria symbiosis-based dating method for Rickettsiales evolution is that it co-estimates the divergence times for both Rickettsiales and their eukaryotic hosts with a unified dataset (Wang and Luo 2021). Notably, under profile mixture models, the origin time of Rickettsiales was estimated at 2.5-2.0 Ga, roughly 0.5 Gyr earlier than that of the last eukaryotic common ancestor (LECA) using homogeneous models estimated in both the current (Fig. 6) and previous studies (Parfrey et al. 2011; Betts et al. 2018; Wang and Luo 2021; Moody et al. 2024). Hence, if the revised time estimate is accurate, it implies that the Rickettsiales’ LCA existed before LECA emerged. This interpretation is consistent with recent comparative genomics findings suggesting that early Rickettsiales initially adopted a free-living or non-obligate symbiotic lifestyle (Castelli et al. 2024), with subsequent evolutionary events leading to independent colonizations of eukaryotic hosts—first protists, then animals (Castelli et al. 2016; Wang and Luo 2021). Alternatively, it is possible that early Rickettsiales were associated with unknown host organisms belonging to eukaryotic stem groups (i.e., extinct or unknown lineages that diverged before LECA but after the split with the closest living relatives of eukaryotes). This hypothesis points to previously unrecognized interactions between stem-group eukaryotes and prokaryotes, and hints at complex prokaryotic engagement as a foundational trait of eukaryotic ancestors, offering insights into the origin of eukaryotes.

## Materials and Methods

### Mixture substitution model

In a mixture model, each category consists of a rate matrix *M*, an equilibrium frequency vector *F*, and a relative substitution rate *R*. Additionally, the computation of the likelihood necessitates the inclusion of branch lengths *θ*, and the tree topology *τ*. Let the full parameter vector be denoted by 𝚯 = (*R*, *F*, *V*, *θ*, *τ*). Denote by ℛ, ℱ, and 𝒱 the number of categories of exchangeability matrices, equilibrium frequencies, and relative substitution rates, respectively. Denote by 𝒮 the number of alignment sites. Hence, the full likelihood of the phylogeny is given as

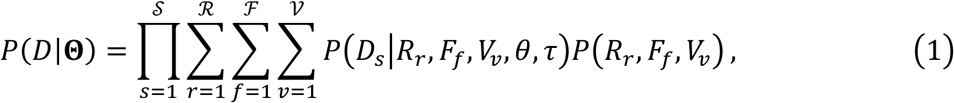

where *P*(*R*_*r*_, *F*_*f*_, *V*_*v*_) indicate the mixture weights (Guindon 2019). Model constraints are enforced by restricting the set of allowed combinations (*r*, *f*, *v*), with the mixture weights normalized over this restricted set. Under independence, this gives *P*(*R*_*r*_, *F*_*f*_, *V*_*v*_) = *P*(*R*_*r*_)*P*(*F*_*f*_)*P*(*V*_*v*_), whereas for linked models such as LG4M or LG4X, combinations with *f* ≠ *v* are excluded.

### Numerical calculation of the Hessian matrix

Two primary numerical methods are commonly employed for computing the Hessian matrix. Let *θ̂* = (*θ̂*_1_, …, *θ̂*_*n*_) indicate the vector of the MLES of branch lengths, and ℓ(*θ̂*) be the log-likelihood of the phylogeny. The outer product of scores (OPS) method (Seo et al. 2004; dos Reis and Yang 2011) estimates the Hessian matrix indirectly using the first derivatives (scores) of the log-likelihood to sum the contributions of the log-likelihood at each site:

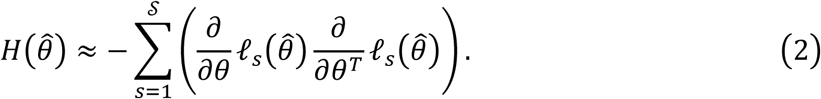

Each component of the score vector 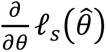 is numerically calculated by the central finite difference method as

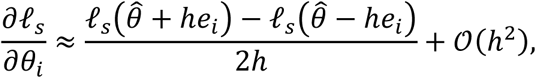

where *e*_*i*_ is the *i*-th standard basis vector, and *h* is a small step size, typically the cube or square root of machine precision.

The second way is by a direct second-order derivative method. It calculates *H*(*θ̂*) via finite differences of the log-likelihood function by

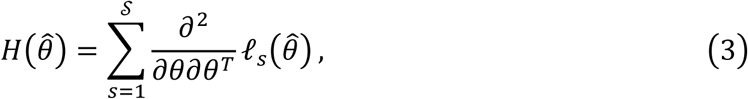

where diagonal elements and off-diagonal elements are respectively calculated by the central finite difference method as

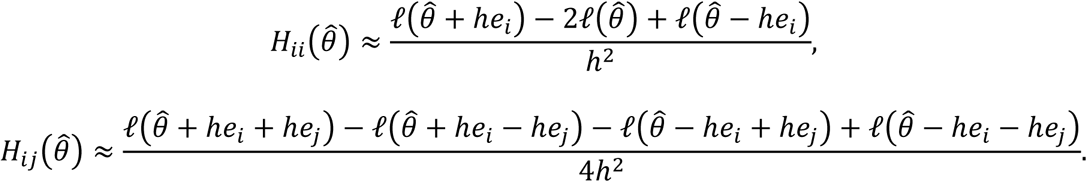

Under regularity conditions, *H*(*θ̂*)/𝒮 converges to the expected Fisher information for a single site 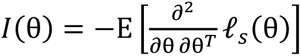 regardless of whether it is calculated using Eq. (2) or (3) (Pawitan 2001):

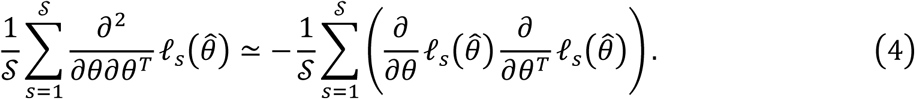

In both methods described as above, the error term is of order 𝒪(*h*^2^). To further improve the numerical accuracy of the second-order derivative method, particularly when dealing with very short branches, we derive the central finite difference calculation of the Hessian at 𝒪(*h*^4^). For the OPS method, this is simply

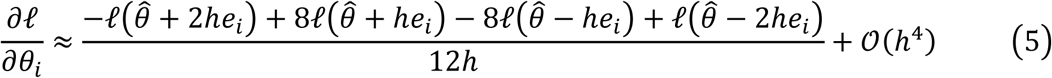

followed by summing the contributions of the log-likelihood at each site according to Eq. (2). Regarding the direct second-order derivative method, the result is more complex and is detailed in Note S3.3.

In phyloHessian, the default algorithm is the OPS method, which is employed by CODEML in PAML v4.3+. Alternatively, users can specify the second-order derivative method, as used in PAML 4.1 and earlier versions. Users can also specify the order of the error term, allowing balancing computational accuracy with speed based on their needs. If not otherwise indicated, the computation of all Hessian matrices in the present study was performed using the OPS algorithm.

### Simulation procedure

We evaluated the performance of different substitution models using simulated data that mimics deep-time evolution (Fig. S16). In our focal simulation scheme, for each substitution model used in sequence evolution simulation (details provided below), we generated 30 timetrees each with 20 tips, using the R package TreeSim (Stadler 2011). Following our previous work (Wang and Luo 2025), timetrees were created under a birth-death model with birth and death rates of 0.4 and 0.2 lineages per 0.1 Gyr as estimated in a prior study of prokaryotic lineage diversification (Scholl and Wiens 2016), and with a taxon sampling proportion of 10% (see Note S1 for alternative settings). Simulated trees were generated with true root ages of 1.0, 2.0, 3.0, and 4.0 Ga to represent a range of evolutionary timescales approaching the age of Earth (∼4.5 Ga).

For each timetree, branch-specific substitution rates were sampled under both the independent rates (IR) and autocorrelated rates (AR) model. For the IR mode, following our previous study (Wang and Luo 2021), the rate follows lognormal distribution with parameters *μ*_IR_ = −3.7 and *σ*_IR_ = 0.2, such that the mean of rate is 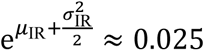 substitutions/site/0.1 Gyr, with a standard deviation of 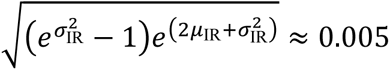 substitutions/site/0.1 Gyr. This roughly corresponds to empirical observations that tip-to-root distances in the bacterial tree of life, constructed with universally conserved orthologs, are roughly 1.0 amino acid substitutions/site (Moody et al. 2022; Wang and Luo 2025), assuming an origin of early life at around 4.0 Ga (Battistuzzi and Hedges 2009; Marin et al. 2017; Betts et al. 2018; Moody et al. 2024). The methodology used to determine the simulation parameters for the AR model is described in detail in the next section.

Alignments of 300 amino acids (aa) were simulated using AliSim (Ly-Trong et al. 2022) according to the timetree generated by simulation, under various substitution models. In the focal analysis, we employed the LG+C60+G4{1.0} model. This indicates LG substitution model plus C60 profile mixture models (60 empirically derived amino acid site frequency profiles) (Quang et al. 2008) and across-site rate variation under a discrete gamma distribution with four relative rate categories parameterized by *⍺* = 1.0, hence Gamma(1,1), meaning that the mean and variance of the across-site relative rates are 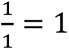 and 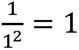 respectively (Yang 1994). Further, we varied several simulation parameters, including sequence length, number of taxa, calibration strategies, birth-death process parameters, across-site rate heterogeneity, molecular clock settings, and model misspecifications. These are detailed in Note S1.

### Determining the parameters for simulation under the AR clock rate model

Denote the absolute rate of sequence evolution (measured in substitutions per site per time unit) under IR and AR respectively by the vectors *X*_IR_ = (*X*_1,IR_, …, *X*_*n*,IR_) and *X*_AR_ = (*X*_1,AR_, …, *X*_*n*,AR_) where *n* represents the number of branches. Their log-transformed values are given by *Y*_IR_ =(*Y*_1,IR_, …, *Y*_*n*,IR_) and *Y*_AR_ = (*Y*_1,AR_, …, *Y*_*n*,AR_). To make simulations under IR and AR models more comparable, we require that the expectations of the sample mean and sample variance of the log-transformed rate under the two models to be equal, respectively, given the same phylogeny. Mathematically, this requires that

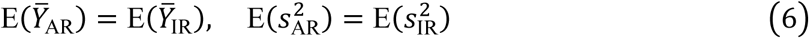

where the sample variances of the log-transformed rates are denoted by 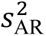 and 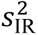.

For the IR model, it is assumed that the rate for each branch, denoted *X*_IR_ = [*X*_1,IR_, …, *X*_*n*,IR_], is independently and identically (*i.i.d.*) distributed according to a lognormal distribution with parameters *μ*_IR_ and 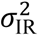, such that the rate for each branch can be expressed as 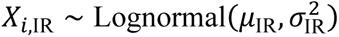. In other words, we have

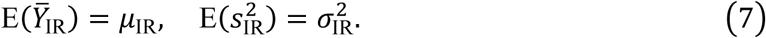

The AR model in molecular clock dating, also known as the geometric Brownian (GBM) model, assumes that the lineage-specific rate follows a geometric Brownian motion with a drift parameter of zero. It is thus parameterized by two parameters, *μ*_AR_, the log-transformed rate at the root, and 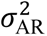, which determines the volatility of the rate changes over time. Accordingly, it can be shown that the log-transformed rate under AR model after time *t* follows 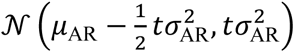 (Rannala and Yang 2007; Panchaksaram et al. 2025). Let 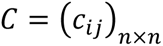 denote the branch-wise covariance matrix where *n* is the number of branches in a rooted tree (note that this is different from what is so-called phylogenetic covariance matrix which is related to only the tips [Fig. S17]). Further, the log-transformed rates *Y*_AR_ follow the following multivariate distribution

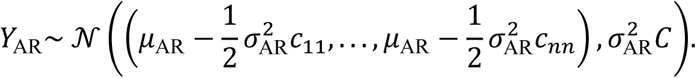

Thus, it is easy to see that

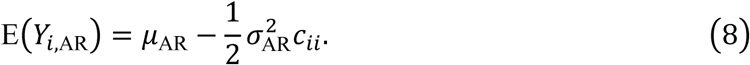

The expectation of the sample variance 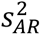 of log-transformed rate 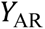 across all *n* branches is calculated as

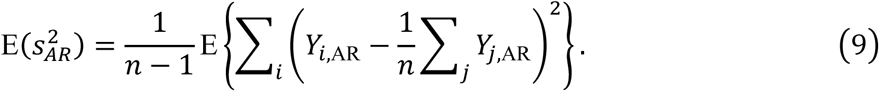

Expanding the right-hand side, we obtain

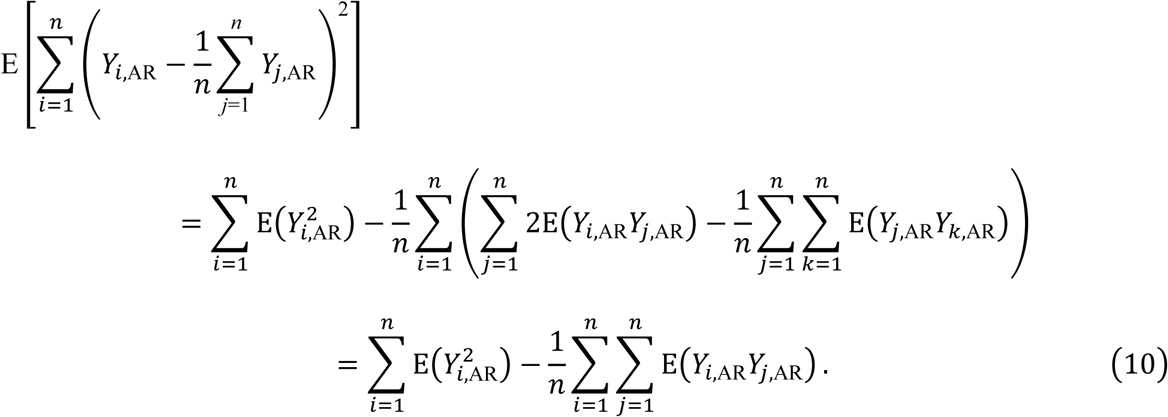

Further, note the following relationship

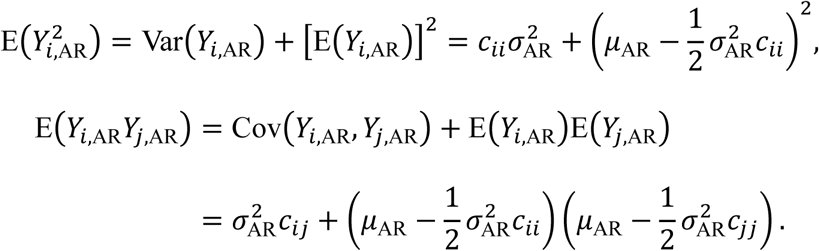

Introducing the above into Eq. (10), we have

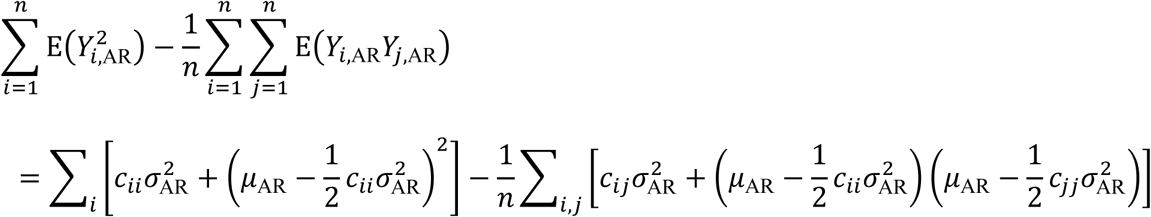

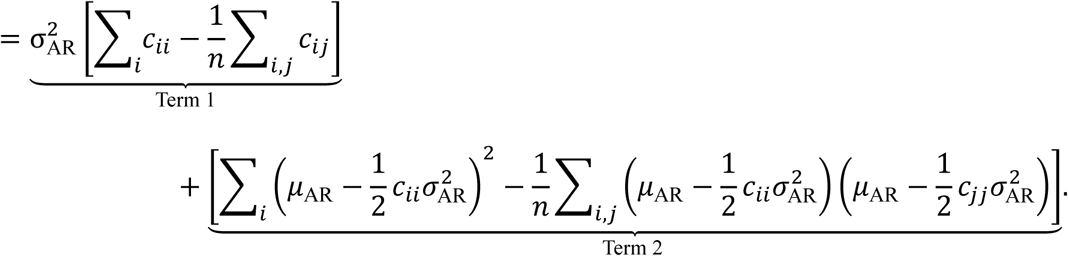

Substituting 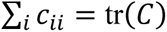 and 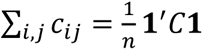 into Term 1, we can rewrite it as

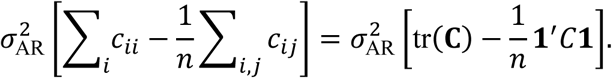

For Term 2, noting that terms involving *μ*_AR_ cancels out, we have

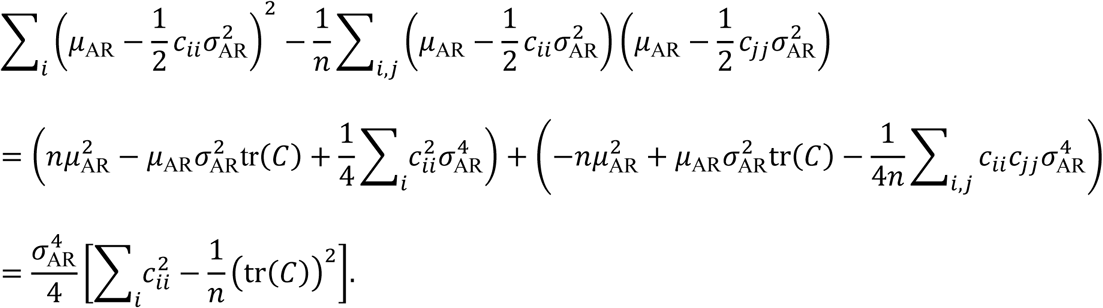

Hence,

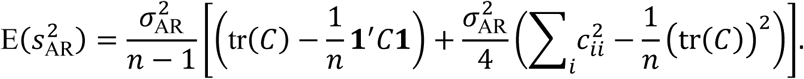

Because 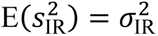, equating 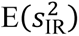 with 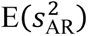 and rearranging terms gives the quadratic equation for 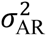

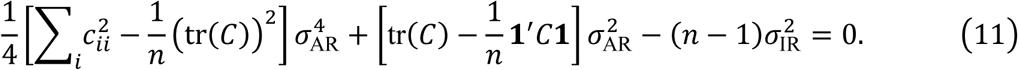

Discarding the negative root, along with solving Eq. (8), we ultimately get

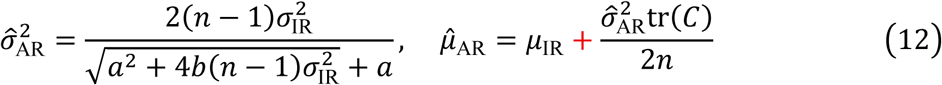

where 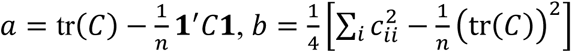 and 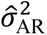 and *μ̂*_AR_ are the values of the parameters for the AR model that satisfy Eq. (6) given the parameters for the IR model (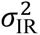 and *μ*_IR_).

### Calibrations used in molecular dating analysis on simulated datasets

Molecular dating analysis were conducted using MCMCtree v4.10.7 (see also Note S3) on phylogenies simulated with procedures described above under varying calibration strategies: a root-only calibration, or root calibration plus one internal calibration, chosen as the node at the median age of all nodes in the true timetree, or two internal calibrations which targeted the 1/3 and 2/3 age quantiles. Time priors for calibrated nodes were set to be uniform within the interval [true_age - (true_age/5), true_age + (true_age/5)], with soft bounds (MCMCtree’s default: 2.5% probability of age outside bounds). Other information about MCMCtree analysis can be found in Note S3. The five calibration schemes mainly used in the present study are *root_only*: a single bounded calibration (both upper and lower bounds); *single_min*: bounded calibration at the root and one minimum-bounded internal node; *single_interval*: bounded calibration at both the root and an internal node; *two_min*: bounded calibration at the root and two minimum-bounded internal nodes; *two_intervals*: bounded calibration at both the root and two internal nodes.

### Comparing the performance of different substitution models on simulated datasets

We evaluated the performance of various substitution models mainly using the relative difference (reldiff) that compared the posterior mean divergence times and substitution rates estimated by MCMCtree (using each model) to the true values used in simulations. Relative difference: calculated as 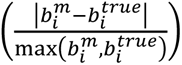, where 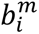 and 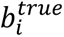 denote the posterior mean divergence times or branch-specific substitution rates inferred by MCMCtree, and those used in simulation (true values), respectively.

### Empirical data analysis

Genomes of Microsporidia and Rickettsiales were downloaded from MicrosporadiaDB (Aurrecoechea et al. 2011) and GenBank (last accessed July 2024), respectively. Sequences of five additional Proteobacteria as the outgroup were retrieved from our prior study (Wang and Luo 2025). TreeCluster v1.0.4 (Balaban et al. 2019) was used to select representative taxa with a cutoff of phylogenetic depth (which takes phylogenetic distance and topology into account to cluster taxa into subclades) of 0.5 and 2.5 respectively for Microsporidia (yielding 27 members) and Rickettsiales (22 members). Genes used to reconstruct phylogeny and molecular dating for Microsporidia were based on 27 orthologs identified by OrthoFinder v3.0.1 (Emms and Kelly 2019) present in all members allowing ≤2 copies per genome, while for Rickettsiales we adopted the mitochondrial endosymbiosis-based strategy (Wang and Luo 2021) by using 24 mitochondria-encoded genes conserved across Alphaproteobacteria (Wang and Wu 2015) (see Data availability). For both datasets, we partitioned the alignments into one, two, or three subsets by fitting the logarithm of the average substitution rates estimated by CODEML into a Gaussian mixture model. We selected the best-fitting substitution models for each partition under both site-homogeneous and site-heterogeneous models according to AIC (Note S3). Calibration information along with alternatives is available in Data S6.

## Data availability

phyloHessian is available at https://github.com/evolbeginner/phyloHessianWrapper. Other codes and the scripts (Goto et al. 2010) generating them, are made available at https://figshare.com/s/96bc117f18051f9642ce.

## Acknowledgements

We particularly thank Minh Bui and Mario dos Reis for comments on early versions of the manuscript and for sharing with us their preprint (Demotte et al. 2025). We also thank Xiyun Jiao, Jianhao Lv, Gergely Szöllősi, Ziheng Yang, and Tianqi Zhu for discussion, Hui Li for proofreading, and Margaret Ip and Haiwei Luo for their support. The work is supported by Natural Science Foundation of China (32400493, 42293294), Hong Kong Research Grants Council (RGC) General Research Fund (GRF) (14112024), and CUHK Direct Grant (4054912).

