## Supplementary material for "Molecular Clock Dating of Deep-Time Evolution Using Complex Mixture Models": SI

### Supplemental Information for “Molecular Clock Dating of Deep-Time Evolution Using Complex Mixture Models”

January 31, 2026

#### Supplemental Figures

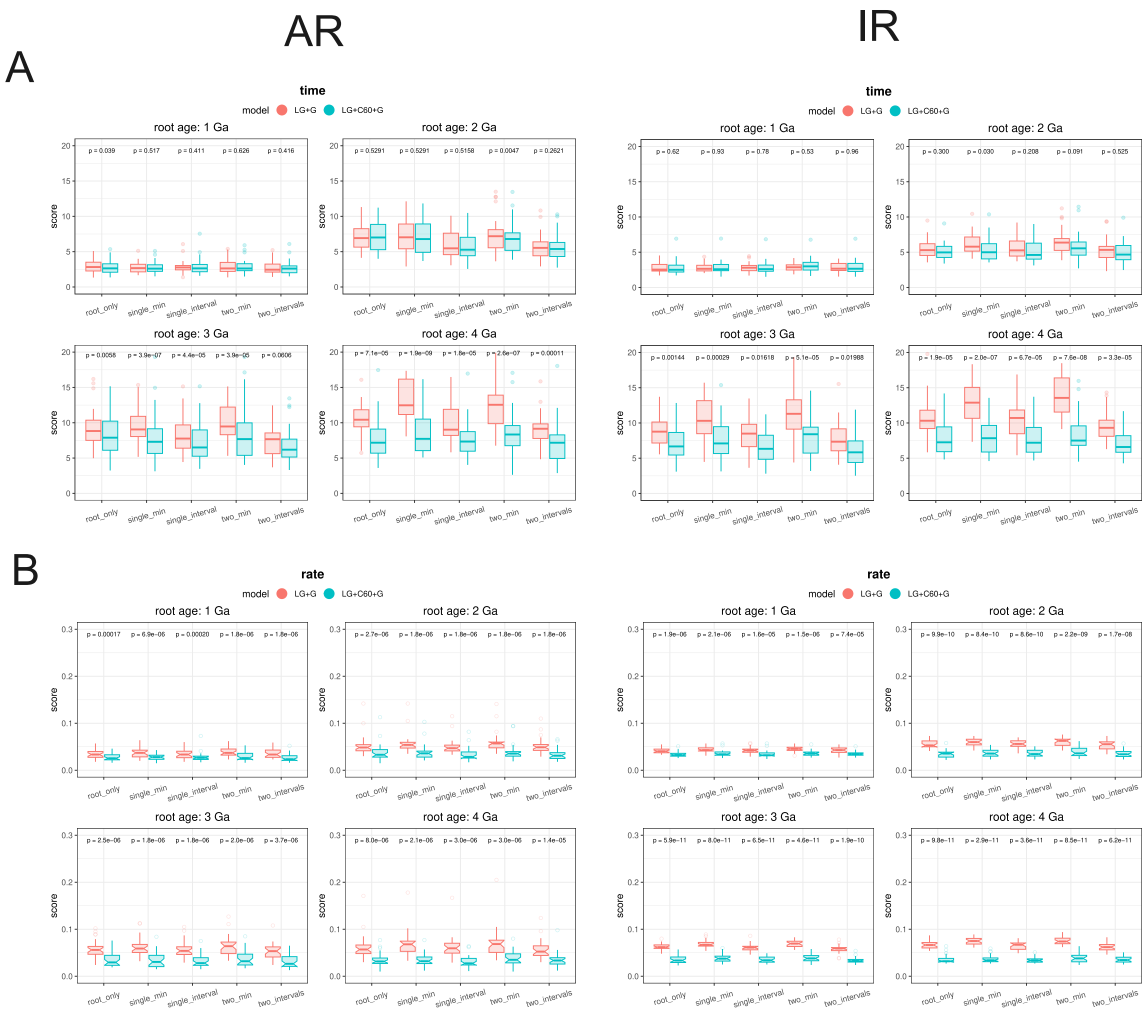


Figure S1: Accuracy of **divergence time** (A) and branch-specific **substitution rate** (B) estimation across simulated root ages (1.0-4.0 Ga). All settings are the same as those used in Fig. 1 except for that the $y$-axis denotes the mean branch score distance (BSD) instead of the relative difference as used in other figures. **BSD** is defined as $\sqrt{\sum_i \left(b_i^m - b_i^{\text{true}}\right)^2}$ where $i$ traverses all branches, and $b_i^m$ and $b_i^{\text{true}}$ denote the posterior ages of branch $i$ in the posterior timetrees under substitution model $m$, and those used in simulation (true values), respectively. Lower values indicate more accurate time estimates under substitution model $m$. Results from autocorrelated-rates (AR) and independent-rates (IR) clock models are displayed on the left and right sides respectively.

A

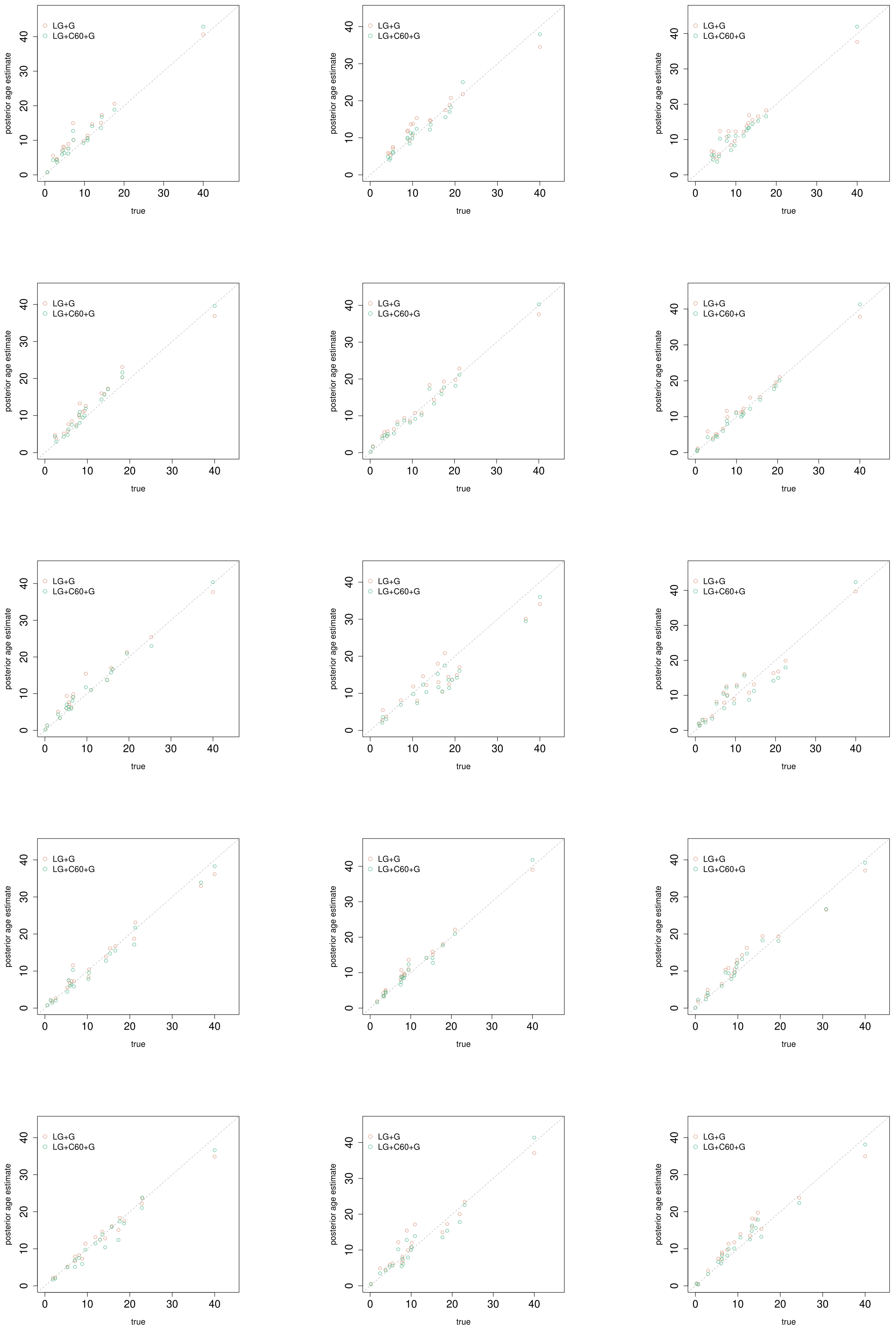

B

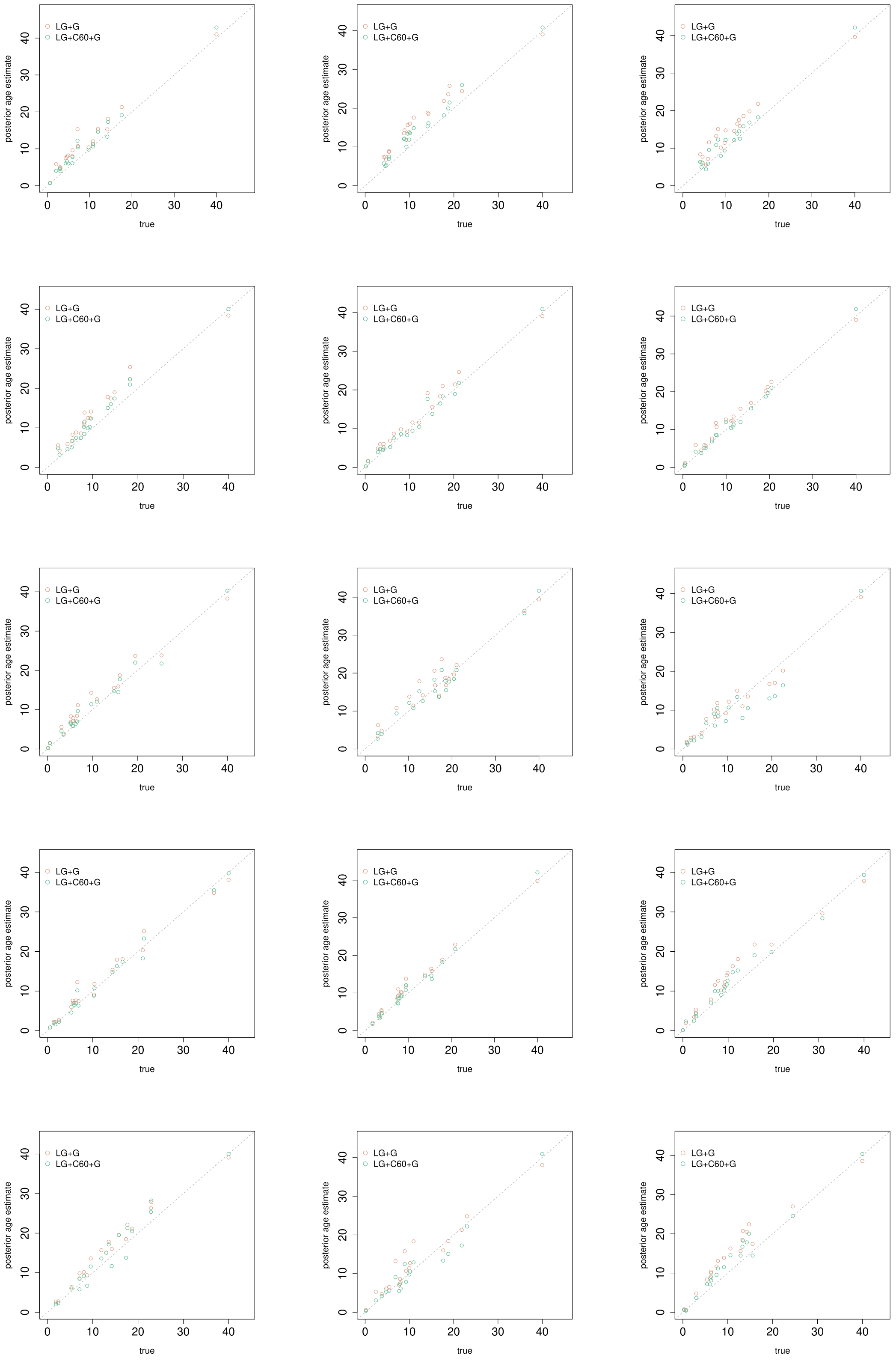


Figure S2: Pairwise comparison of the posterior mean **divergence times** between LG+G (*x*-axis) and LG+C60+G (*y*-axis) under the **AR** (autocorrelated-rates) clock-rate model in simulation. This represents a visualization of the results corresponding to the calibration strategies *root_only* (A) and *two_min* (B), respectively, in Fig. 1 with the root age in simulation of 4.0 Ga. Only the first 15 of the 30 simulation results are displayed for simplicity. The time unit is 100 million years (Myr).

A

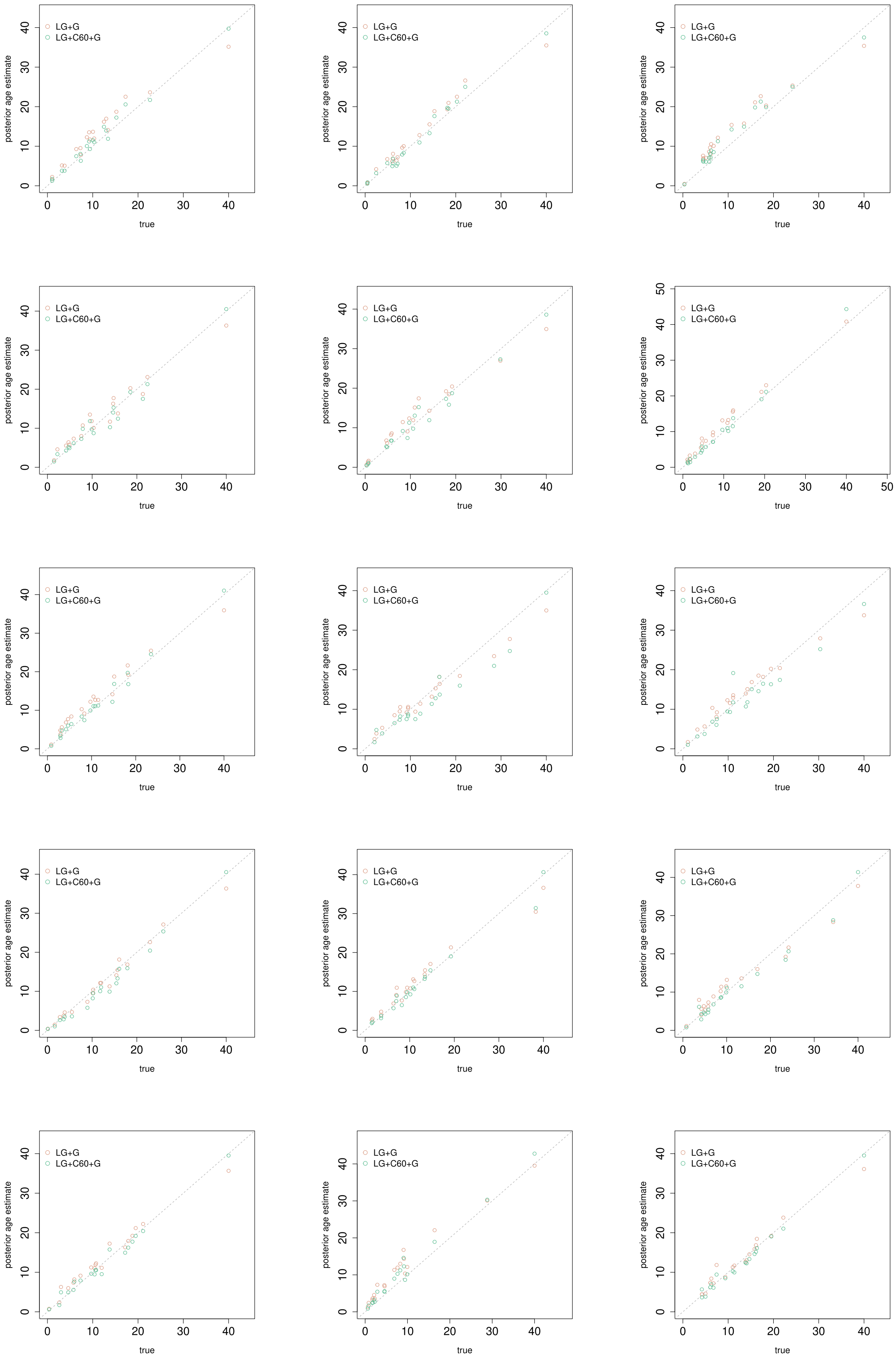

B

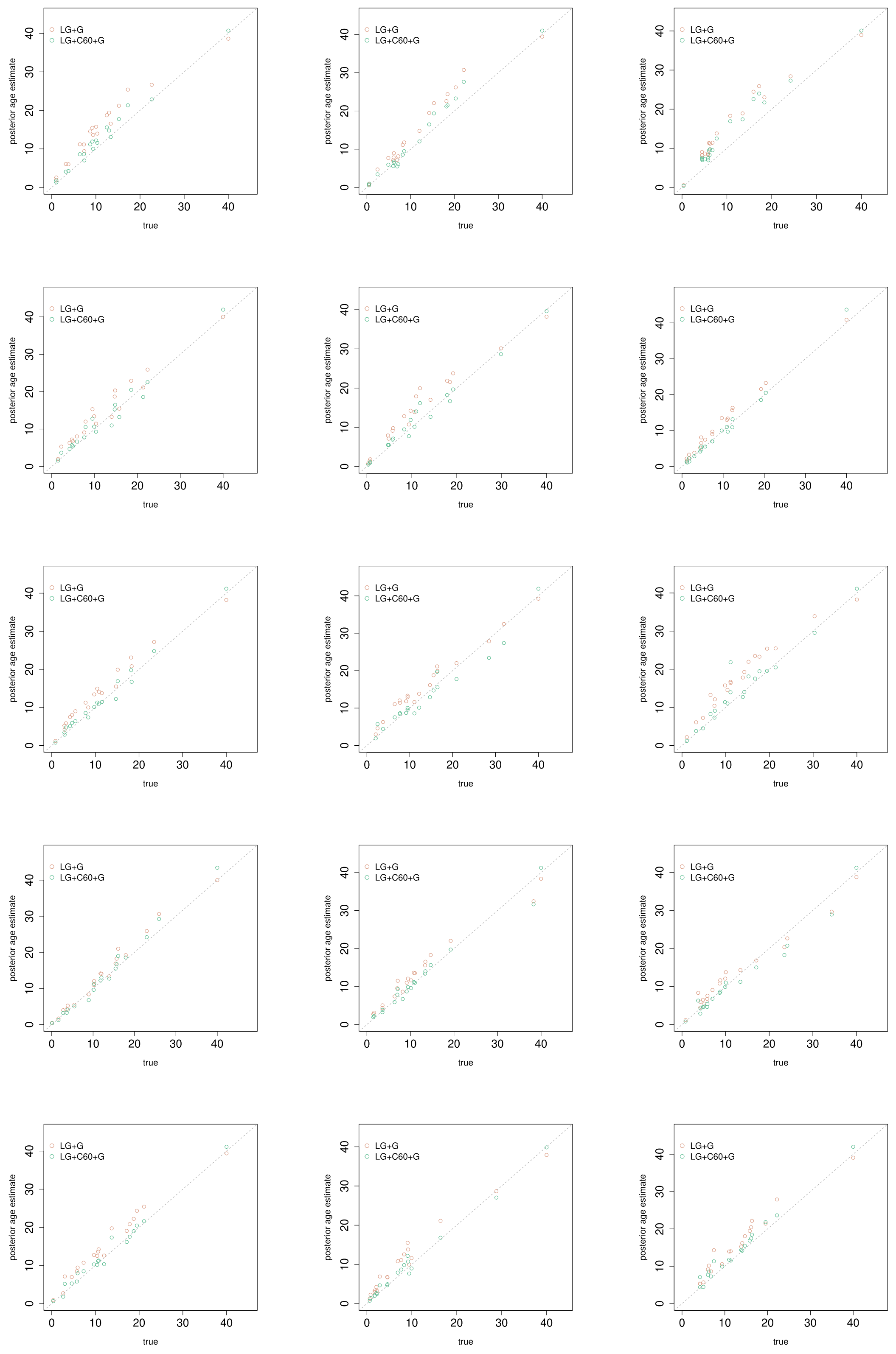


Figure S3: Pairwise comparison of the posterior mean **divergence times** between LG+G (*x*-axis) and LG+C60+G (*y*-axis) under the **IR** (independent-rates) clock-rate model in simulation. This represents a visualization of the results corresponding to the calibration strategies *root_only* (A) and *two_min* (B), respectively, in Fig. 1 with the root age in simulation of 4.0 Ga. Only the first 15 of the 30 simulation results are displayed for simplicity. The time unit is 100 million years (Myr).

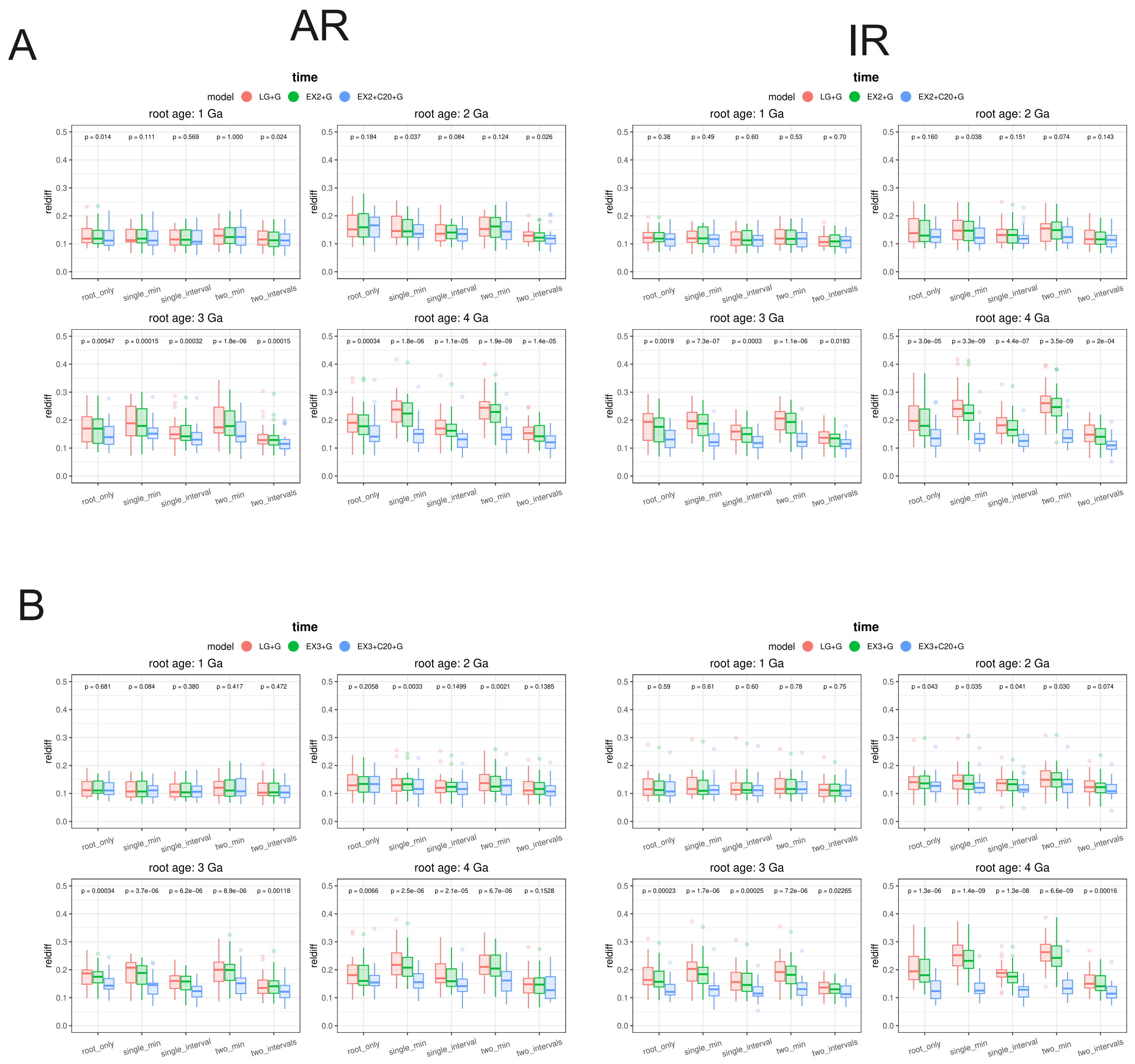


Figure S4: Accuracy of **divergence time** estimation under simulated mixture models EX2+C20+G (A) and EX3+C20+G (B). Root ages are set as 1.0, 2.0, 3.0, 4.0 Ga. Each boxplot represents the average of the relative difference compared to the simulated value from 30 simulations. Calibration strategies *root_only*, *single_min*, *two_min*, *single_interval*, *two_intervals* are the same as Fig. 1. *P*-values are derived from a paired Wilconxon test between the LG+G model and the true (mixture) model. Results from AR and IR clock models are displayed on the left and right sides respectively.

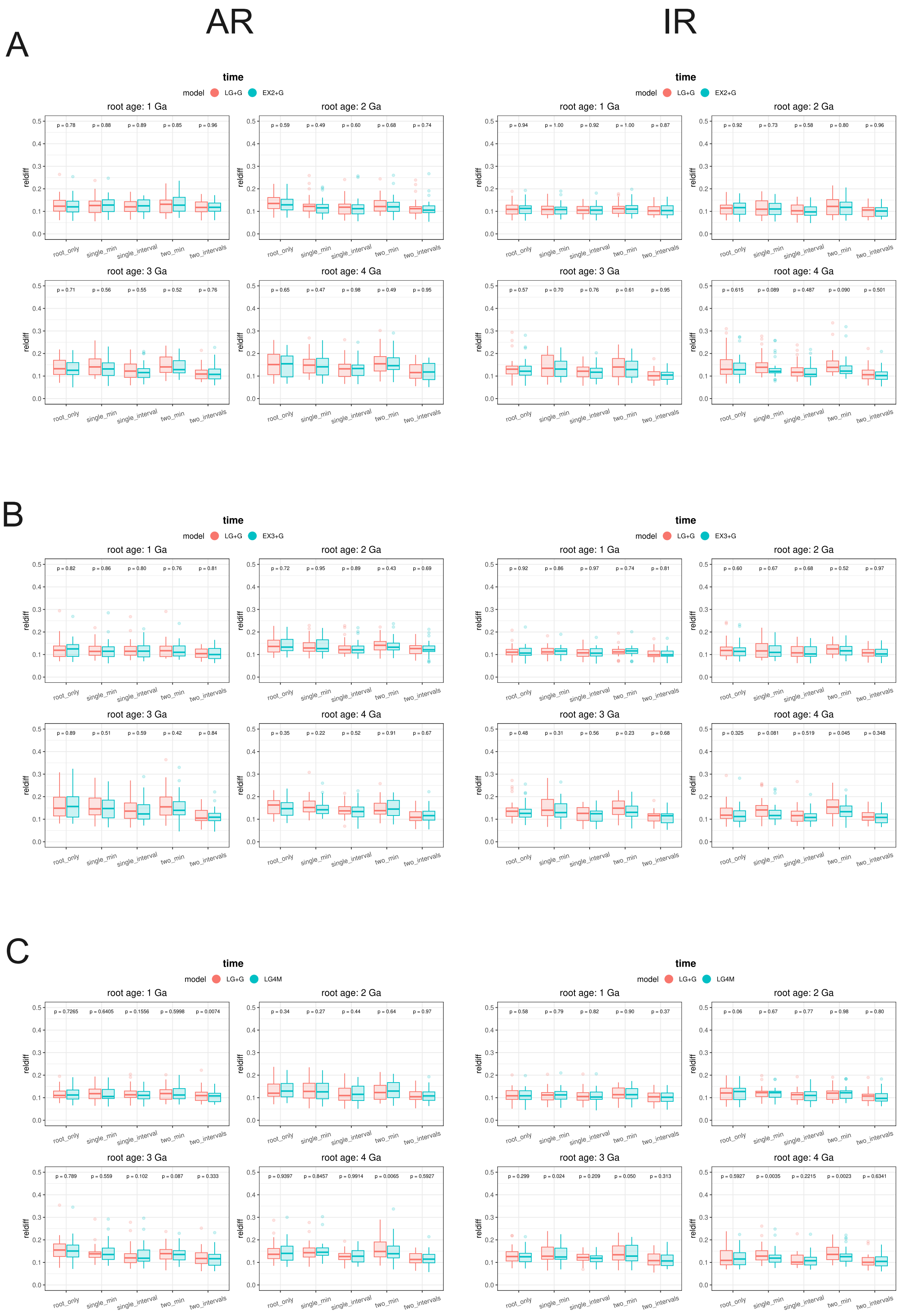


Figure S5: Accuracy of **divergence time** estimation under simulated mixture models EX2+G (A), EX3+G (B), and LG4M (C). Root ages are set as 1.0, 2.0, 3.0, 4.0 Ga. Each boxplot represents the average of the relative difference compared to the simulated value from 30 simulations. Calibration strategies *root_only*, *single_min*, *two_min*, *single_interval*, *two_intervals* are the same as Fig. 1. *P*-values are derived from a paired Wilconxon test between the LG+G model and the true (mixture) model. Results from AR and IR clock models are displayed on the left and right sides respectively.

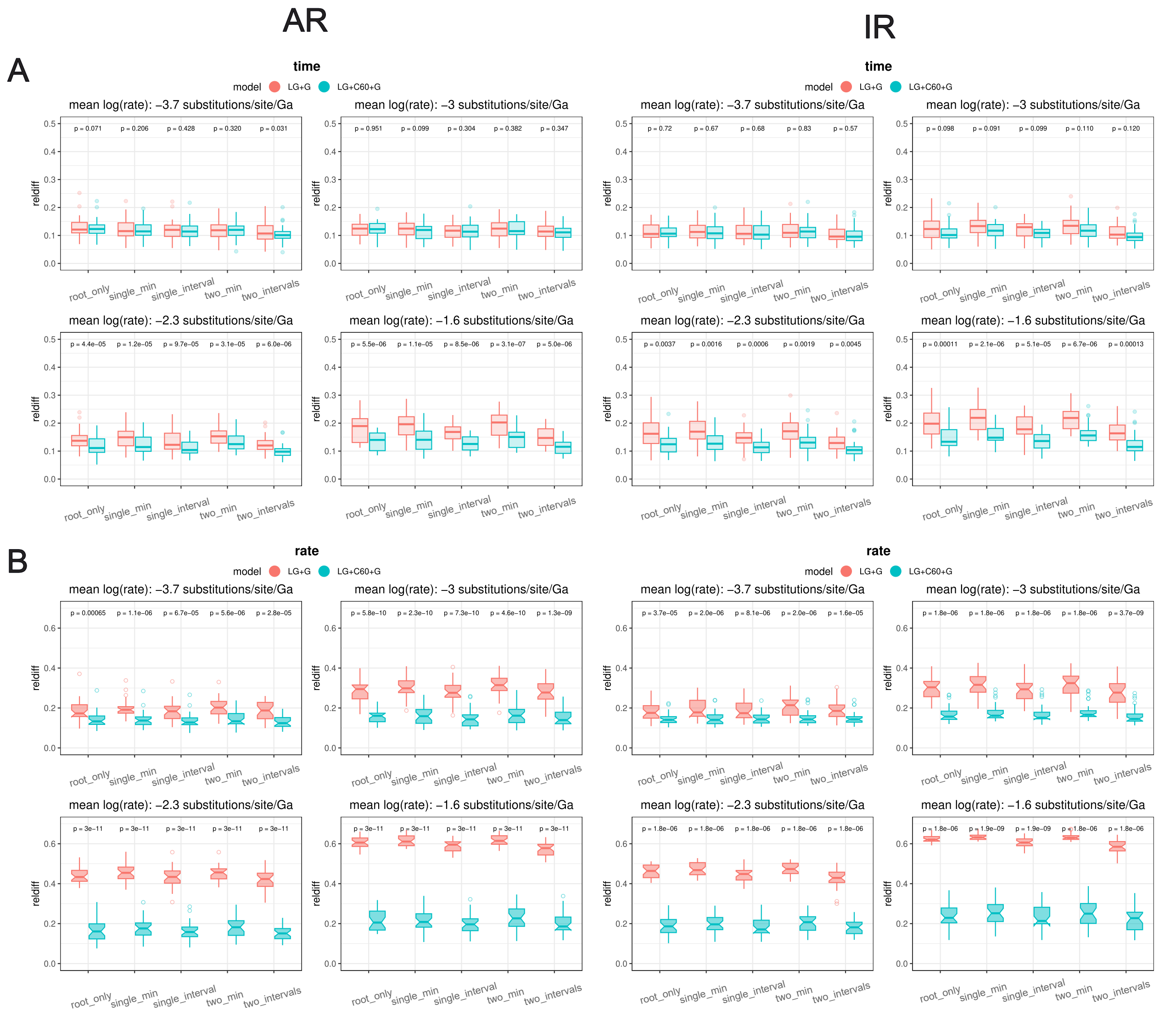


Figure S6: Accuracy of **divergence time** estimation with **varying substitution rates**. For each panel, we generate 30 timetrees (each with 20 tips) with a fixed root age of 1.0 Ga. The substitution rates vary (0.025, 0.05, 0.075, 0.1 substitutions/site/0.1 Gyr). This was achieved for the IR model, by setting lognormal distribution parameter $\mu$ to -3.7, -3.0, -2.3, and -1.6 substitutions/site/0.1 Gyr while keeping $\sigma = 0.2$, and for the AR model, by setting the parameters such that the expectation and variance of the log-rate satisfy Eq. (12) in the main text (see also Materials and Methods). The $y$-axis represents the relative difference. Calibration strategies are identical to those described in Fig. 1.

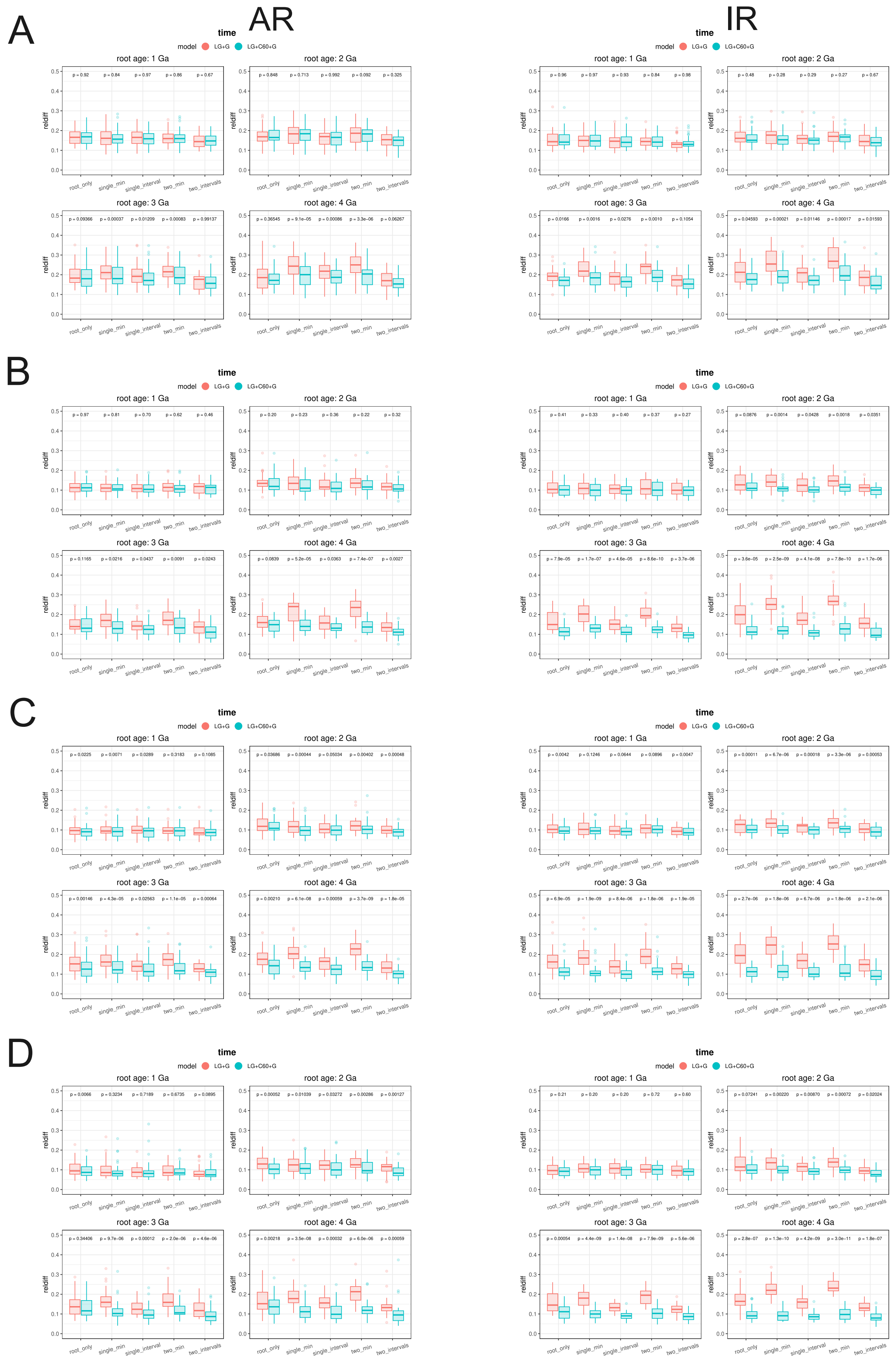

Figure S7: Accuracy of **divergence time** estimation with different sequence lengths. Alignments of varying lengths (100 [A], 500 [B], 1000 [C], and 2000 [D] amino acids) are simulated using the LG+C60+G{1} substitution model. Clock models AR (left) and IR (right) are applied, with root ages ranging from 1.0 to 4.0 Ga. The y-axis represents the relative difference. All other settings are the same as Fig. 1

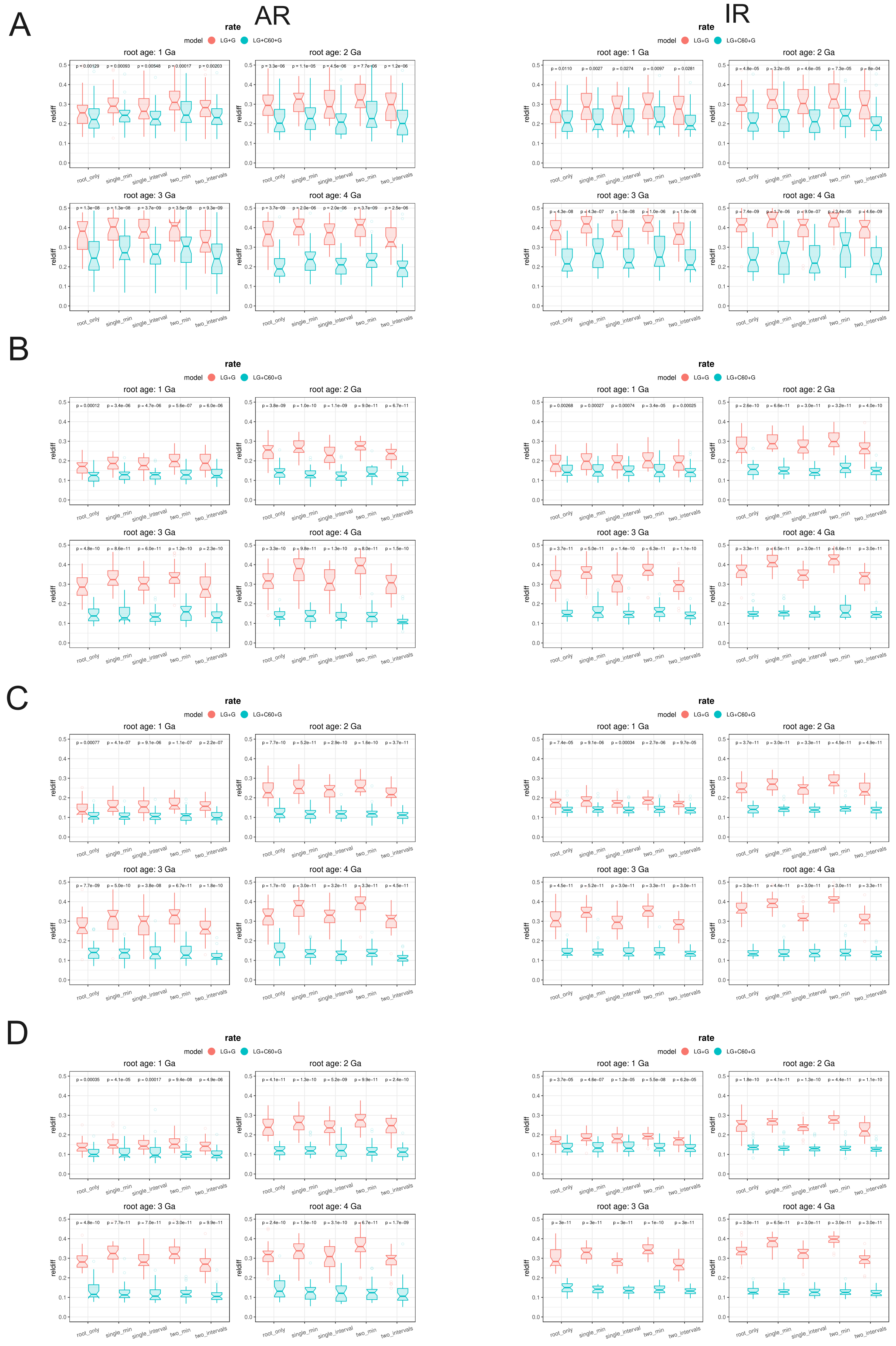


Figure S8: Accuracy of branch-specific **substitution rate** estimation with different sequence lengths. Alignments of varying lengths (100 [A], 500 [B], 1000 [C], and 2000 [D] amino acids) are simulated using the LG+C60+G4{1.0} substitution model. Clock models AR (left) and IR (right) are applied, with root ages ranging from 1.0 to 4.0 Ga. The y-axis represents the relative difference. All other settings are the same as Fig. 1.

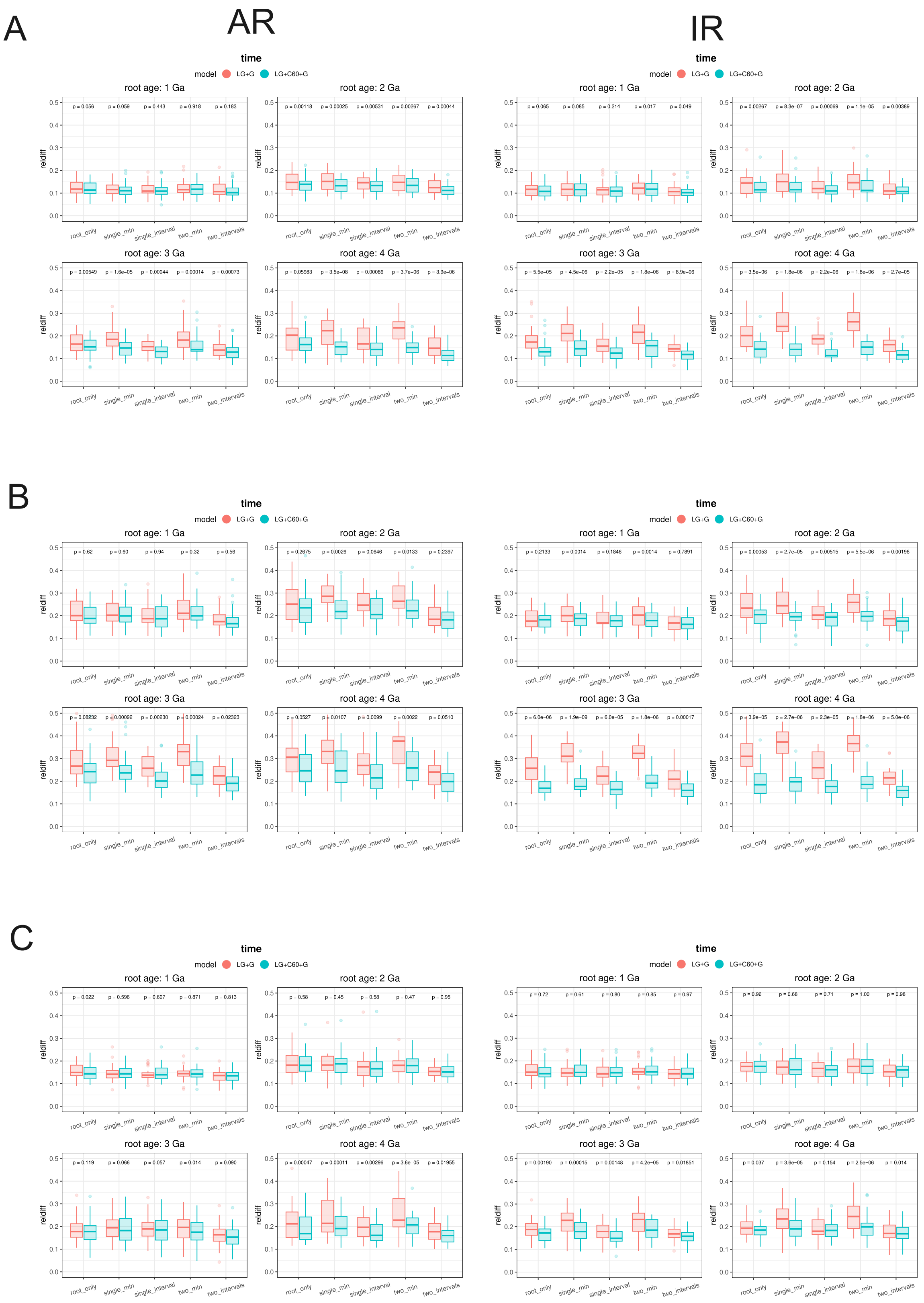


Figure S9: Accuracy of **divergence time** estimation under alternative settings. (A) *Gamma_1.5*: LG+C60+G{1.5} is used in simulation. (B) *complete_taxon_sampling*: birth-death prior with birth rate ($\lambda$) = 0.4 lineages per 0.1 Gyr, death rate ($\mu$) = 0.2 lineages per 0.1 Gyr, and sampling proportion ($\rho$) = 0.1. (C) *SD_lograte_0.55*: a log-normal distribution for the branch-specific rate, with $\sigma = 0.55$, implying greater across-branch rate variation. See Note S1 for more detailed information for each alternative simulation scheme. Root ages are set as 1.0, 2.0, 3.0, 4.0 Ga. Each boxplot represents the average of the relative difference compared to the simulated value from 30 simulations. Calibration strategies *root_only*, *single_min*, *two_min*, *single_interval*, *two_intervals* are the same as Fig. 1. The sequence length is 300 aa (if not otherwise indicated).

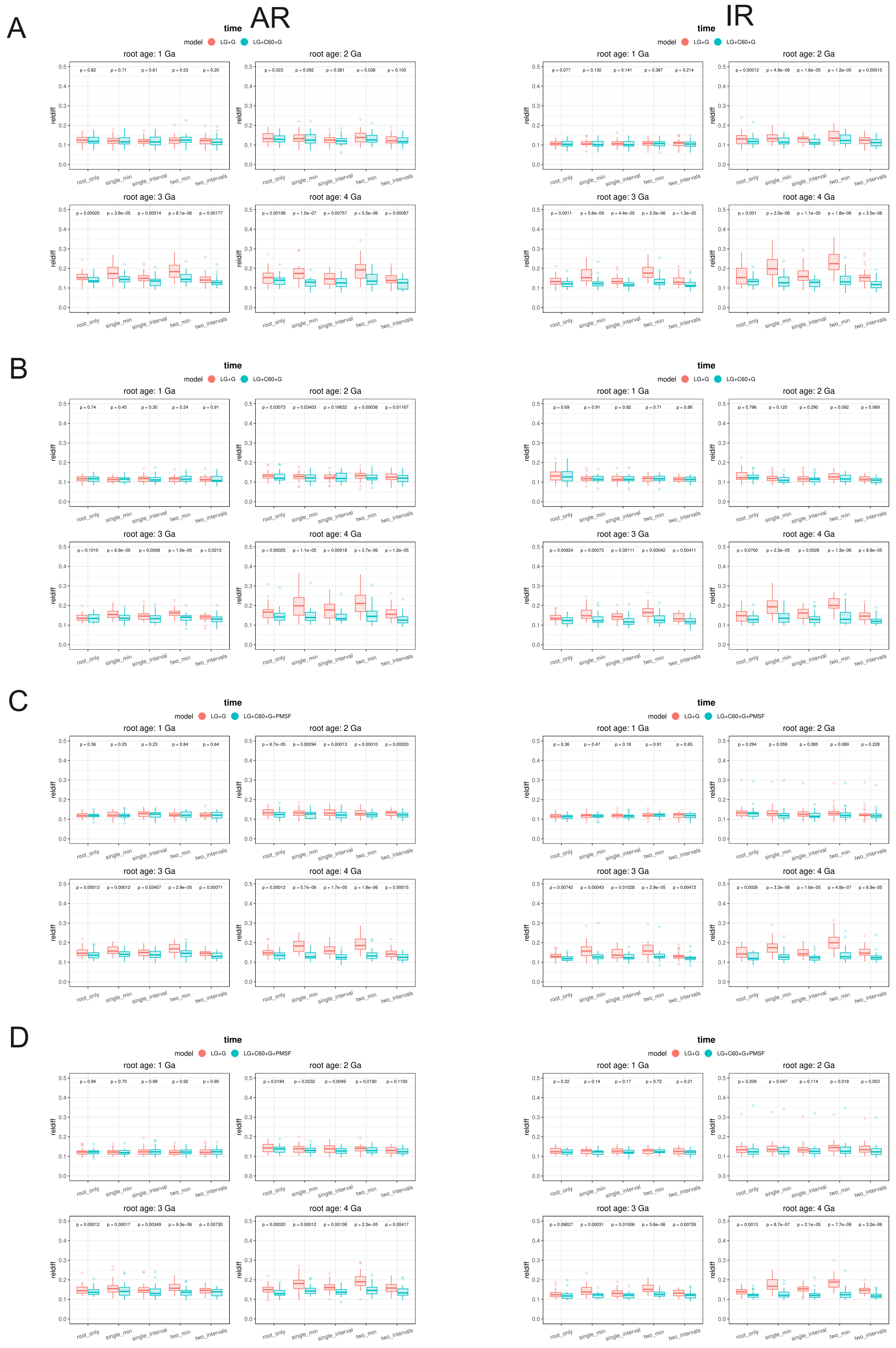


Figure S10: Accuracy of **divergence time** estimation with different tip numbers. Alignments of 40 [A], 60 [B], 80 [C], and 100 [D] tips are simulated using the LG+C60+G4{1.0} substitution model. Clock models AR (left) and IR (right) are applied, with root ages ranging from 1.0 to 4.0 Ga. The *y*-axis represents the relative difference. All other settings are the same as Fig. 1. Mixture model dating analyses with 80 (C) or 100 tips (D) are performed under the PMSF approximation (LG+C60+G+PMSF) to reduce computational time.

A

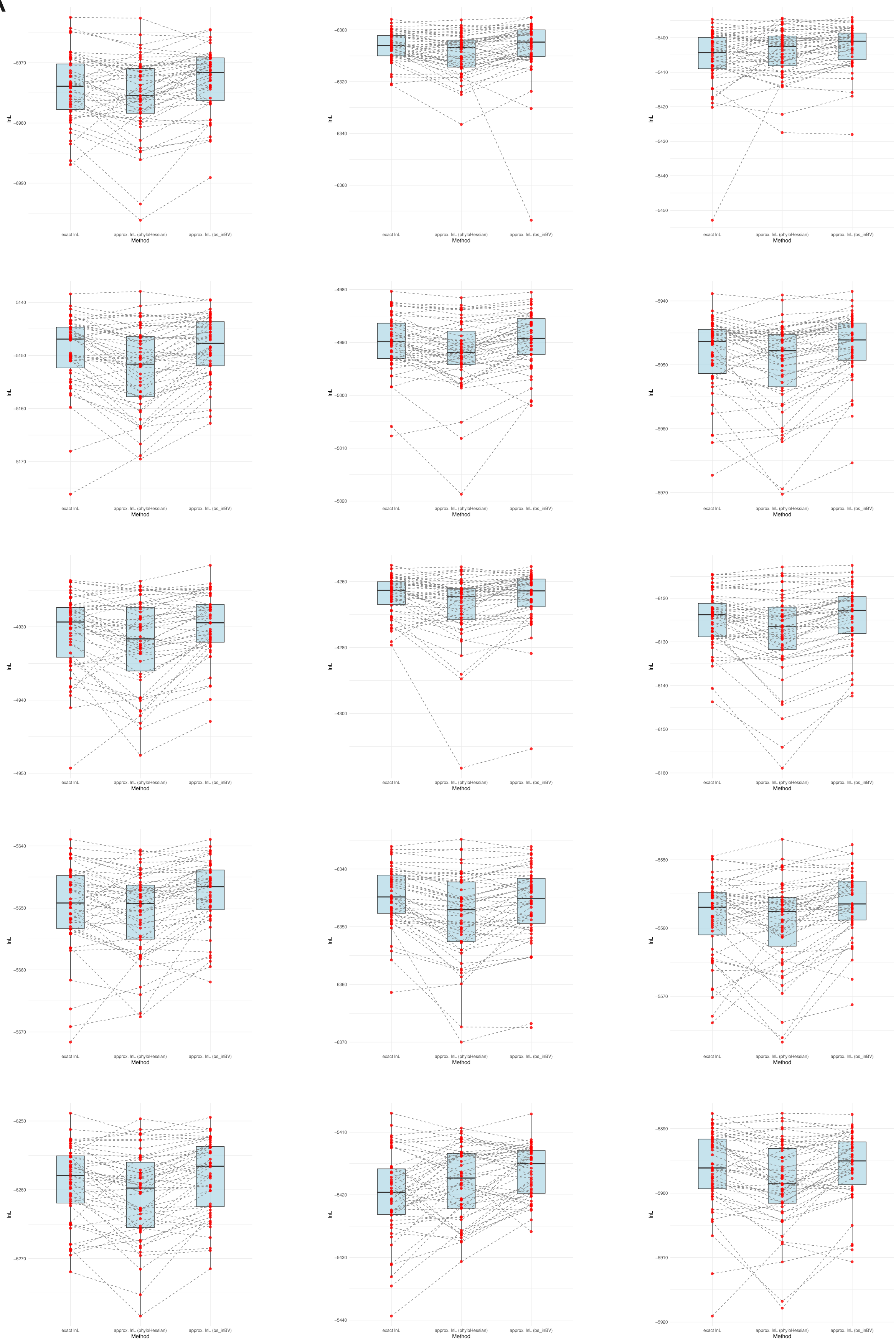
lnL
Method
exact lnL
approx. lnL (phyloHessian)
approx. lnL (bs_inBV)

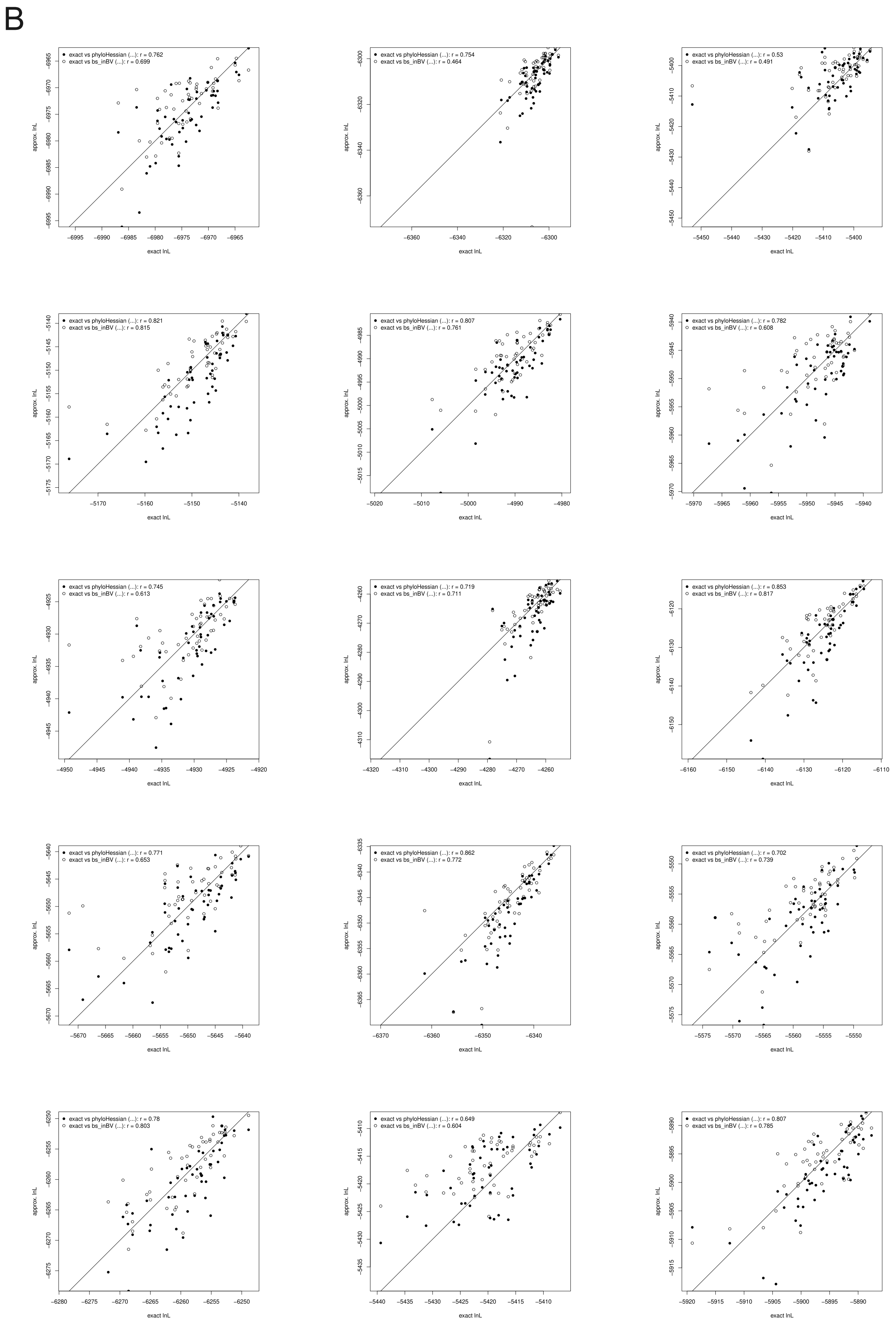


Figure S11: Comparison of the log-likelihoods (lnL) calculated with the exact method, approximate likelihood method by phyloHessian, and approximate likelihood method by bs_inBV (bootstrap) under LG+C20+G (for simplicity). lnLs are calculated with branch lengths from the first 50 out of 1000 Felsenstein's bootstrap (Felsenstein, 1985) alignments. (A) Paired-boxplot comparison of the lnLs calculated with the exact method (left), approximate likelihood method by phyloHessian (finite difference; middle), and approximate likelihood method by bs_inBV (bootstrap; right). (B) Scatter plot with $x-$axis representing the exact lnL, and $y-$axis the approximated lnL estimated by phyloHessian (filled dots) and bs_inBV (open dots). $r$ indicates the Pearson correlation coefficient. No transform (Reis and Yang, 2011) is applied to the branch lengths.

A

Divergence time (unit: 0.1 Gyr) AR

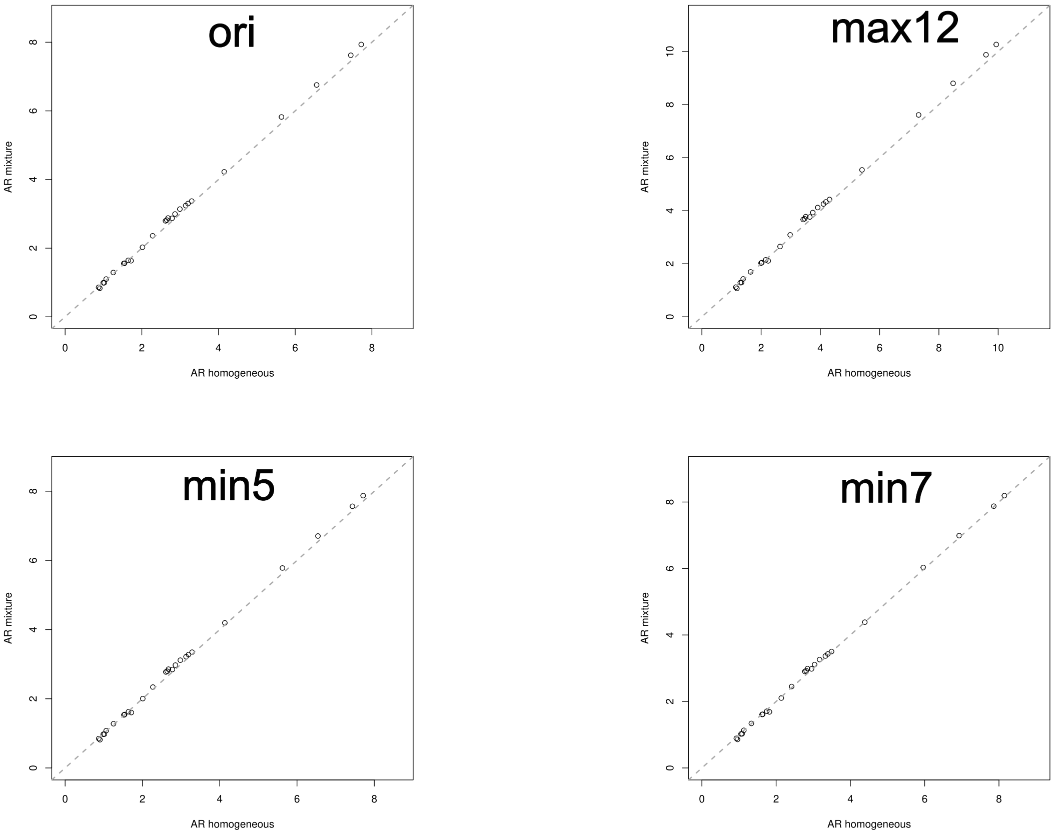


B

Divergence time (unit: 0.1 Gyr) IR

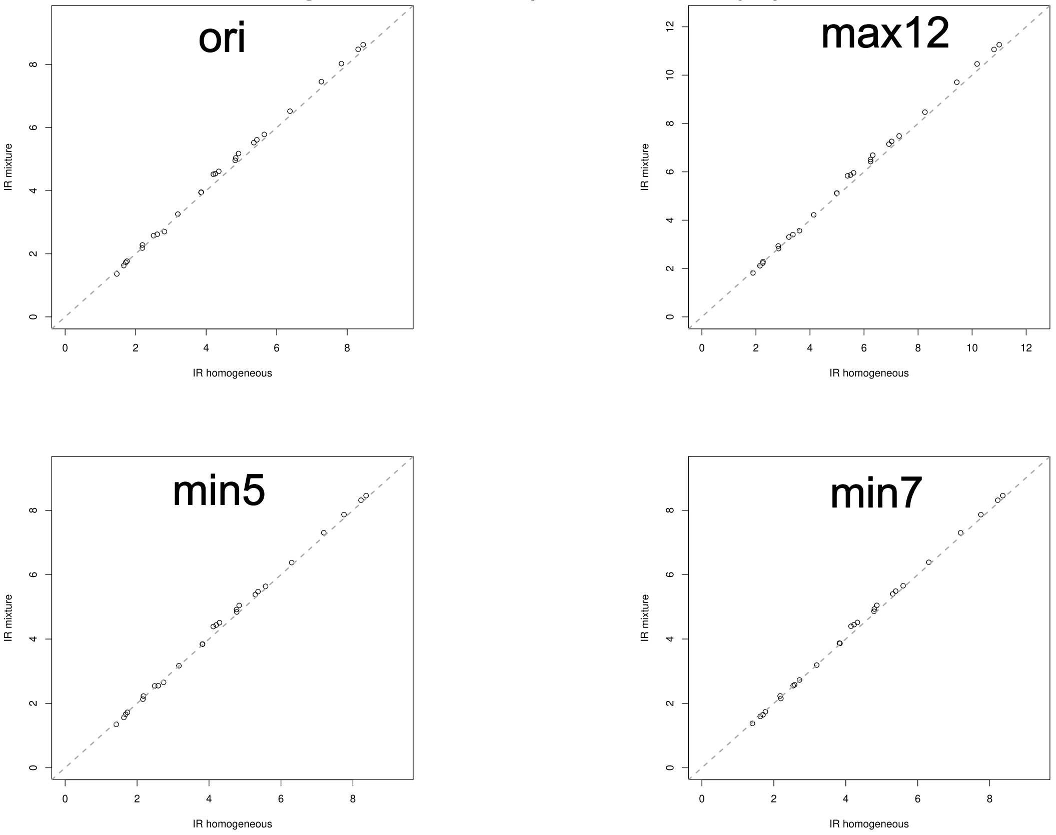

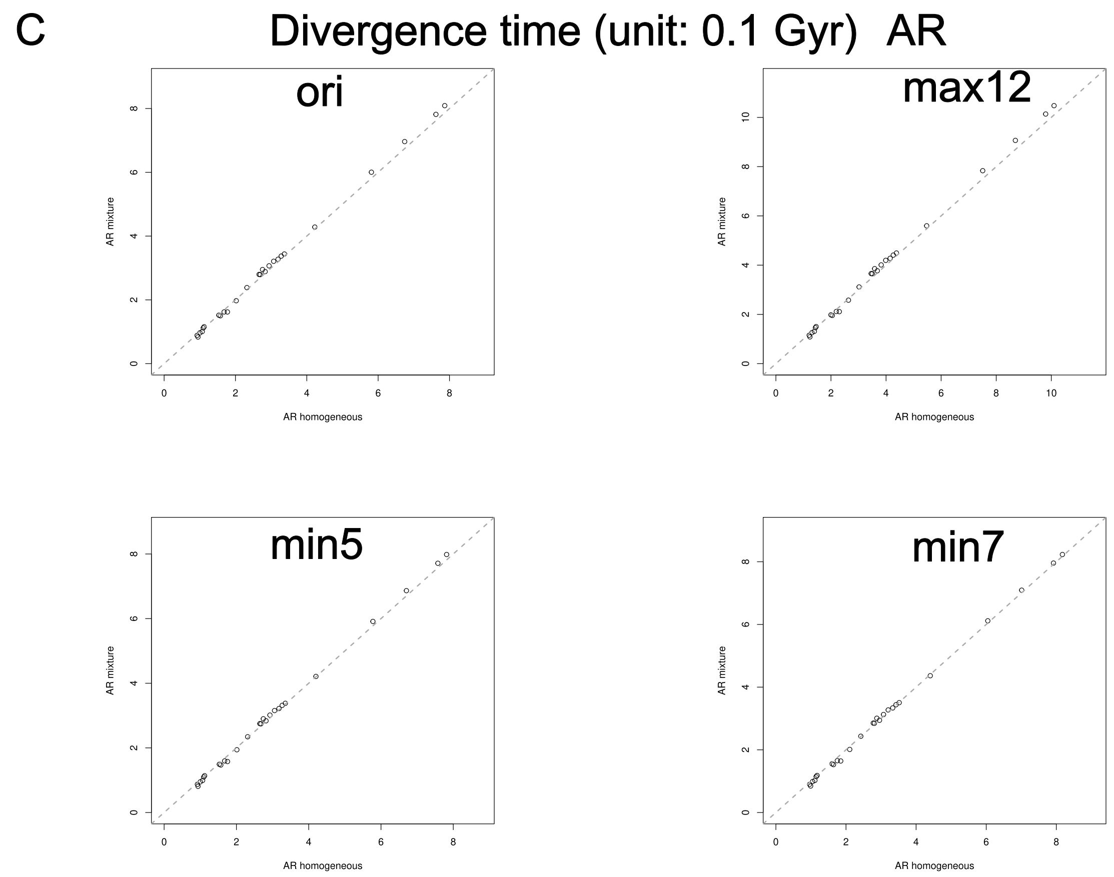
C
Divergence time (unit: 0.1 Gyr) AR
ori
max12
min5
min7
AR mixture
AR homogeneous

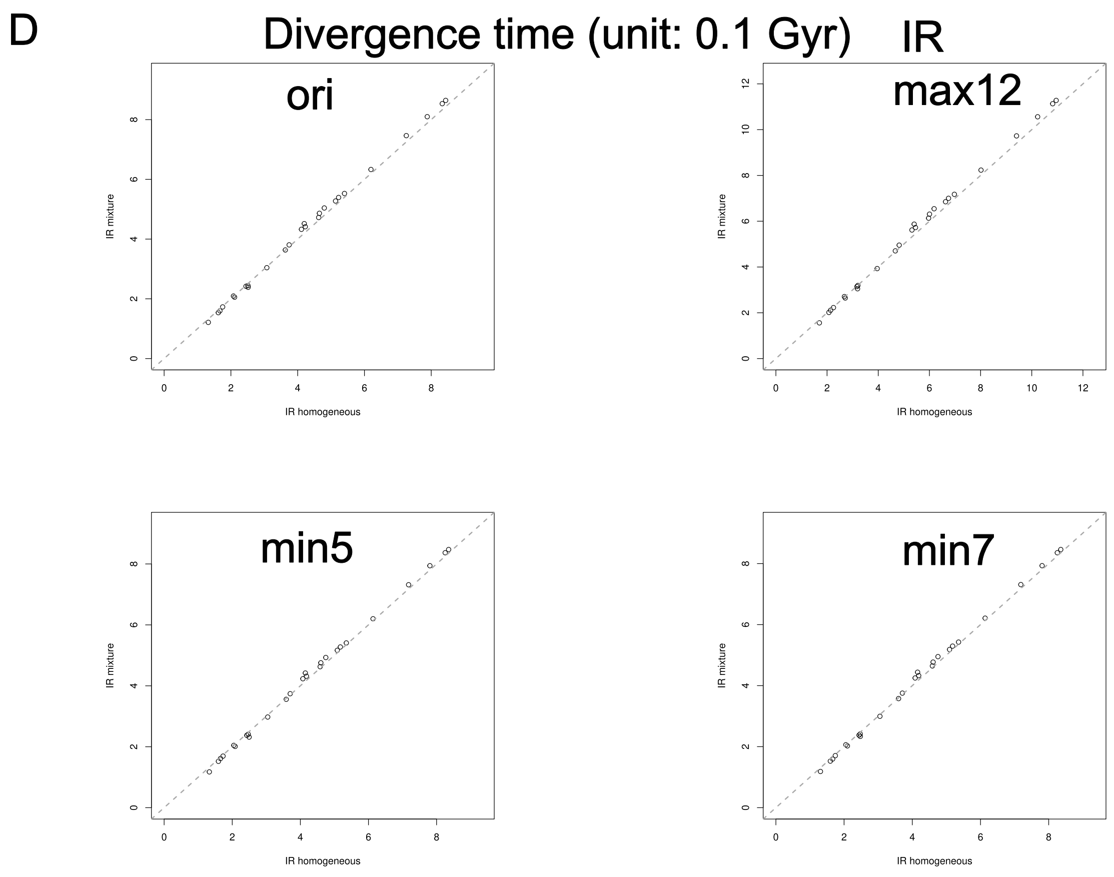
D
Divergence time (unit: 0.1 Gyr) IR
ori
max12
min5
min7
IR mixture
IR homogeneous

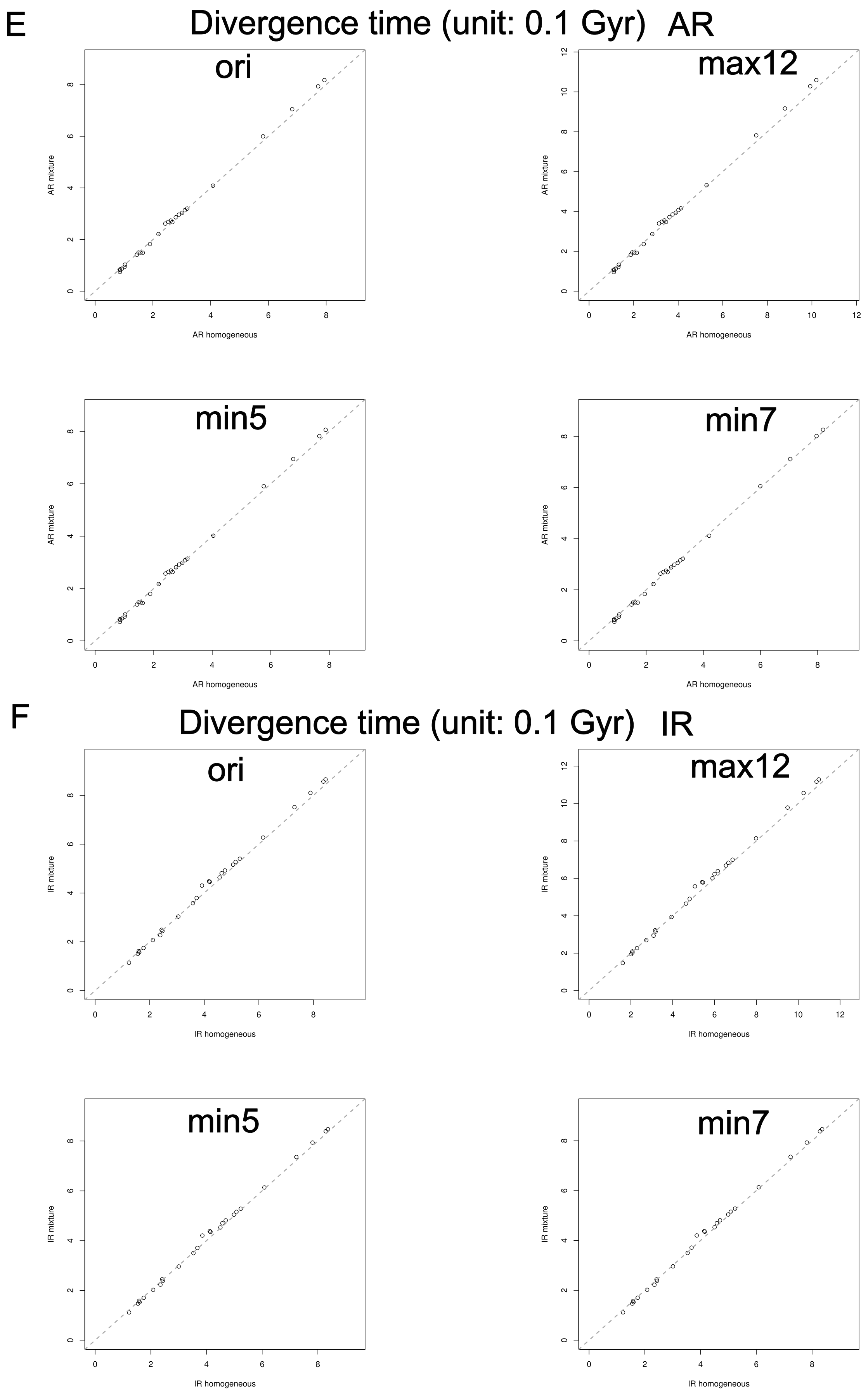


Figure S12: Comparison of estimated divergence time between the best-fitting homogeneous ($x$-axis) and mixture models ($y$-axis) with four different calibration strategies for **Microsporidia**. (A) A single partition under the AR clock model. (B) A single partition under the IR clock model. (C) Two partitions under the AR clock model. (D) Two partitions under the IR clock model. (E) Three partitions under the AR clock model. (F) Three partitions under the IR clock model. Sequence alignment is partitioned into different numbers of subsets according to their estimated substitution rates using a Gaussian mixture model on the log-transformed values (see Methods). Details of the different calibration strategies can be found in Data S6.

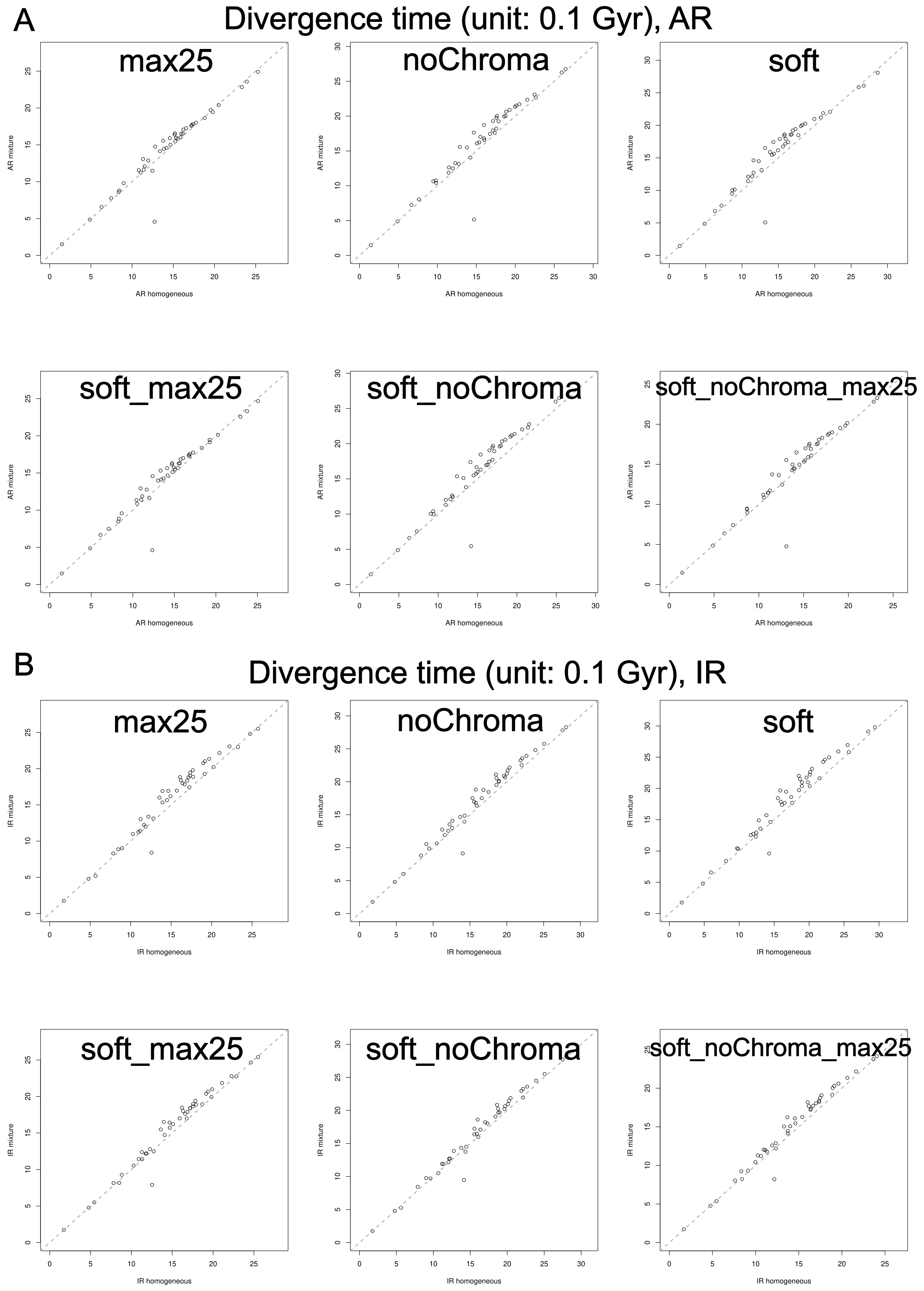

A
Divergence time (unit: 0.1 Gyr), AR
max25
noChroma
soft
soft_max25
soft_noChroma
soft_noChroma_max25
AR mixture
AR homogeneous
B
Divergence time (unit: 0.1 Gyr), IR
max25
noChroma
soft
soft_max25
soft_noChroma
soft_noChroma_max25
IR mixture
IR homogeneous

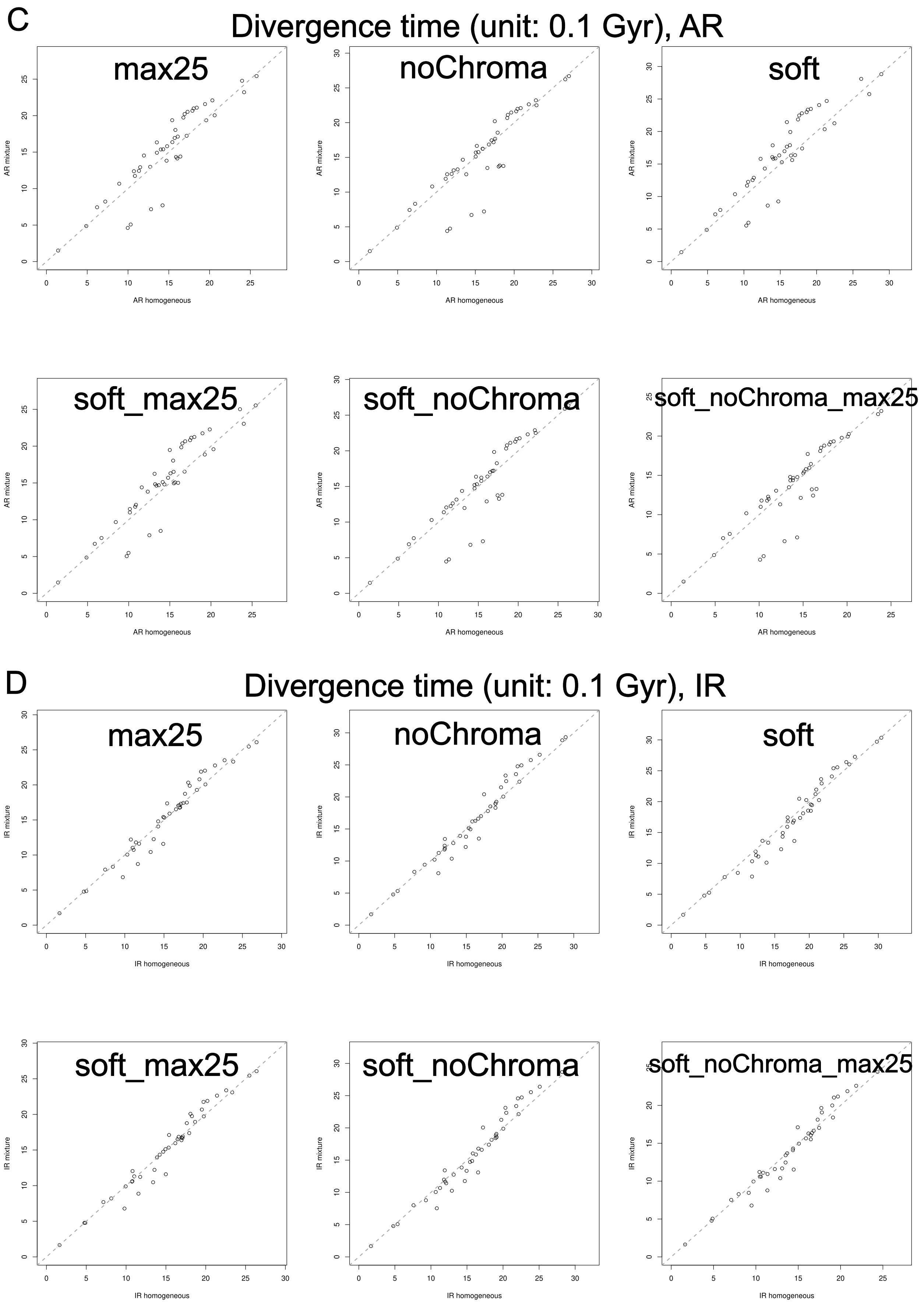
C
Divergence time (unit: 0.1 Gyr), AR
max25
noChroma
soft
soft_max25
soft_noChroma
soft_noChroma_max25
AR mixture
AR homogeneous
D
Divergence time (unit: 0.1 Gyr), IR
max25
noChroma
soft
soft_max25
soft_noChroma
soft_noChroma_max25
IR mixture
IR homogeneous

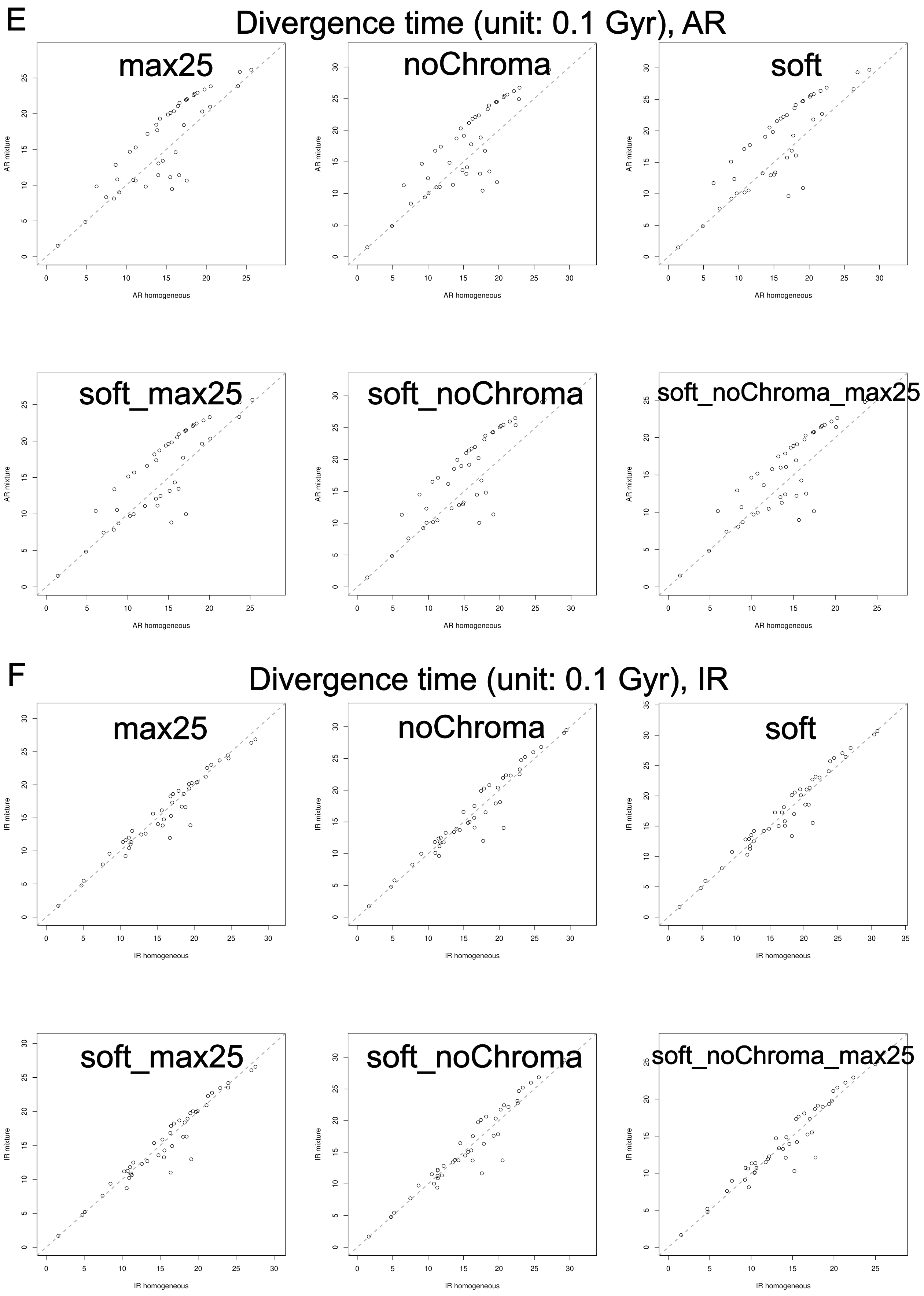


Figure S13: Comparison of posterior mean divergence time between the best-fitting homogeneous ($x$-axis) and mixture models ($y$-axis) with eight different calibration strategies for **Rickettsiales**. (A) A single partition under the AR clock model. (B) A single partition under the IR clock model. (C) Two partitions under the AR clock model. (D) Two partitions under the IR clock model. (E) Three partitions under the AR clock model. (F) Three partitions under the IR clock model. Sequence alignment is partitioned into different numbers of subsets according to their estimated substitution rates using a Gaussian mixture model on the log-transformed values (see Methods). Details of the different calibration strategies can be found in Data S6.

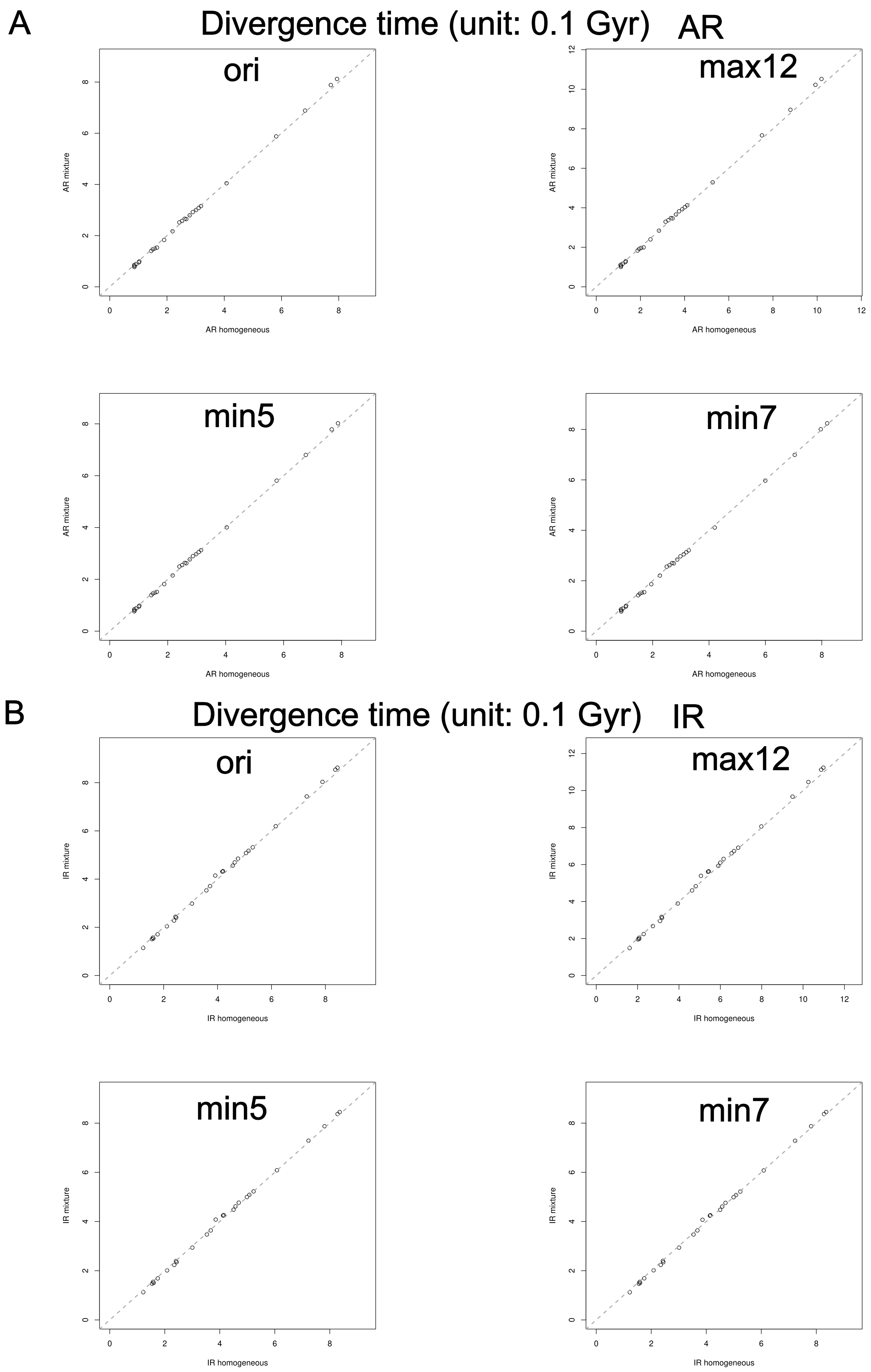


Figure S14: Comparison of posterior mean divergence time between the best-fitting homogeneous ($x$-axis) and mixture models ($y$-axis) with four different calibration strategies for **Microsporidia** with best-fitting models **without the +F (empirical frequency)** option for the mixture models. (A) Three partitions and AR clock model. (B) Three partitions and IR clock model. Details of the different calibration strategies can be found in Data S6.

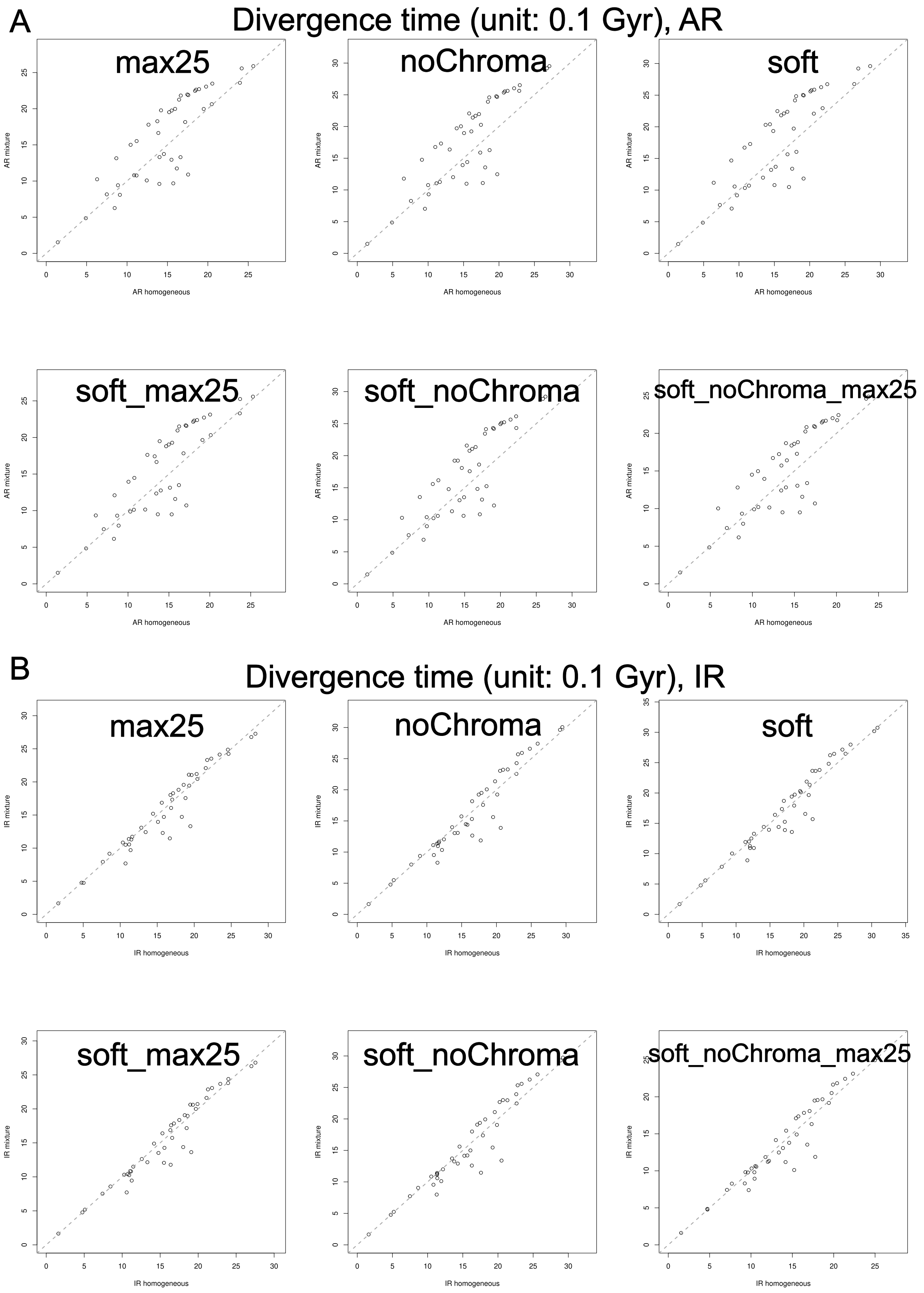


Figure S15: Comparison of divergence time and substitution rate between the best-fitting homogeneous ($x$-axis) and mixture models ($y$-axis) with eight different calibration strategies for **Rickettsiales** with best-fitting models **without the +F (empirical frequency)** option for the mixture models. (A) Three partitions and AR clock model. (B) Three partitions and IR clock model. Details of the different calibration strategies can be found in Data S6.

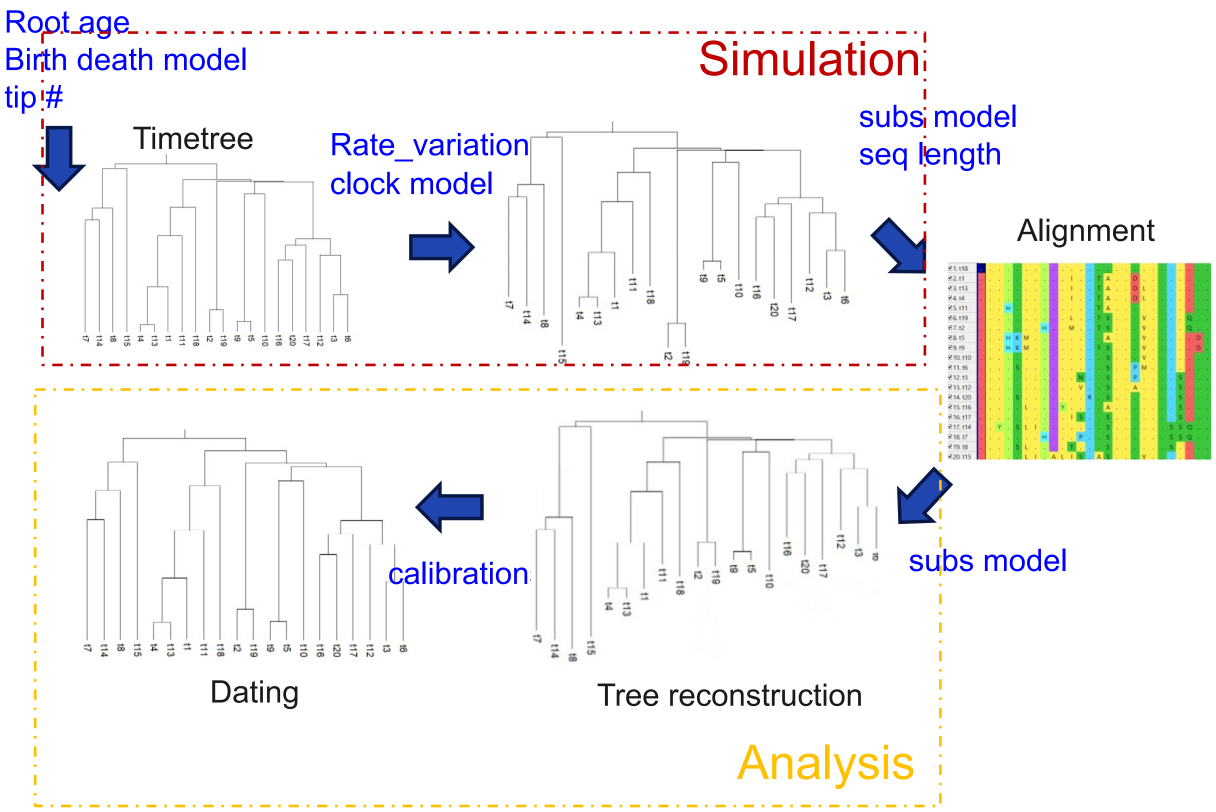


Figure S16: A schematic figure showing the procedure of simulation to study the impact of different substitution models in molecular clock analysis. Blue texts indicate where alternative settings are tested (Note S1).

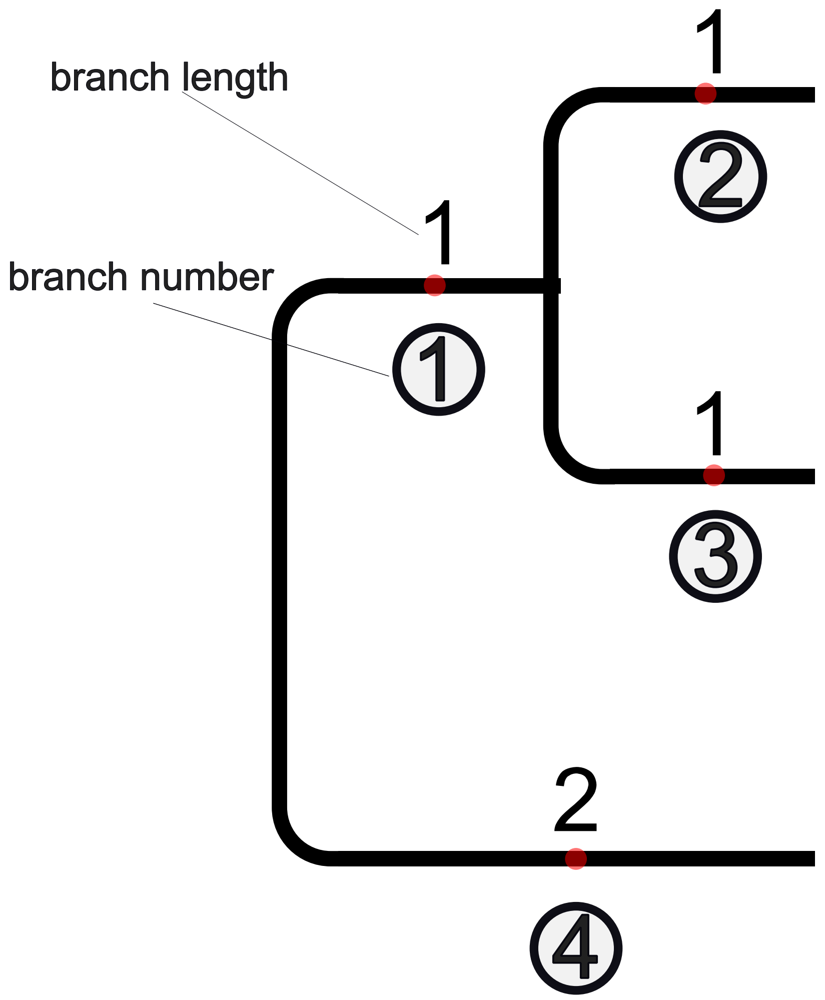


branchwise covariance matrix $C$

$$\begin{array}{c c} & \begin{array}{cccc} ① & ② & ③ & ④ \end{array} \\ \begin{array}{c} ① \\ ② \\ ③ \\ ④ \end{array} & \begin{pmatrix} 0.5 & 0.5 & 0.5 & 0 \\ 0.5 & 1.5 & 1.0 & 0 \\ 0.5 & 1.0 & 1.5 & 0 \\ 0 & 0 & 0 & 1 \end{pmatrix} \end{array}$$

Figure S17: An illustration of the branch-wise phylogenetic variance-covariance matrix. The length of each branch is given above the branch. The circled numbers serve as labels for the branches, positioned below each branch. Note that the distance is measured at the midpoint of each branch (denoted by a red circle), consistent with the AR molecular clock model's simplification of the mean substitution rate of the branch to the rate at the midpoint. The branch-wise phylogenetic variance-covariance matrix for this working example is shown at the bottom of the figure.

### Note S1. Simulation schemes

*1) Focal*

Details of the focal simulation scheme are given in Materials and Methods in the main text. For an easy comparison across schemes in Supplemental Data, we briefly describe it as follows. A number of 30 timetrees each with 20 tips were generated using TreeSim (Stadler 2011). Timetrees were created under a birth-death model with birth and death rates of 0.4 and 0.2 lineages per 0.1 Gyr respectively, and the taxon sampling proportion of 10%. Simulated trees were generated with true root ages of 1.0, 2.0, 3.0, and 4.0 Ga to represent a range of evolutionary timescales approaching the age of Earth (~4.5 Ga). Branch-specific substitution rates were sampled under both IR and AR clock models. For the IR mode, the mean rate of each branch follows a lognormal distribution with the parameters $\mu_{\text{IR}} = -3.7$ and $\sigma_{\text{IR}} = 0.2$ (such that $\frac{\text{sd(rate)}}{\text{mean(rate)}} \approx \frac{0.005}{0.025} = 0.2$), while the AR model parameters were chosen to match the expectations of the sample mean and sample variance of the branch-wise rates of the IR model. Alignments of 300 amino acids (aa) were simulated using AliSim according to the timetree generated by simulation. In the focal analysis, we applied LG+C60+G4{1.0} (or simply LG+C60+G where unambiguous; "+G4{1.0}" specifies a discrete Gamma distribution of four rate categories with the shape parameter $\alpha = 1.0$) (figs. 1-3).

*2) Diff_rate*

Root age was fixed at 1.0 Ga, while mean substitution rates varied (0.025, 0.05, 0.075, 0.1 substitutions/site/0.1 Gyr) by setting lognormal distribution parameter $\mu$ to -3.7, -3.0, -2.3, and -1.6 substitutions/site/0.1 Gyr, representing a range from normal to very fast evolution. This contrasts with the focal scheme where the root age varied from 1.0 to 4.0 Ga and the mean of the substitution rates was fixed at 0.025 substitutions/site/0.1 Gyr (fig. S6).

#### Birth-death model

*3) flat_time_prior*

A nearly flat (uniform) prior to the ages of internal nodes. The birth-death process is parameterized by birth and death rates of 1 and 1 lineages per 0.1 Gyr and taxon sampling proportion equal to 0.001 (fig. 2B).

*4) complete_taxon_sampling*

Ages of internal nodes from birth-death process of a non-flat prior. The birth-death process is parameterized by birth and death rates of 0.4 and 0.2 lineages per 0.1 Gyr and taxon sampling frequency equal of 1.0 (fig. S9B).

#### Across-branch rate variance

*5) SD_lograte_0.38*

Identical to the focal scheme, but with the variance of the logarithm of rate equal to $0.38^2$ instead of $0.2^2$, such that $\frac{\text{sd(rate)}}{\text{mean(rate)}} = \frac{\sqrt{\left(e^{0.2^2}-1\right)e^{(-2\times 3.7+0.38^2)}}}{e^{-3.7+\frac{0.38^2}{2}}} \approx 0.4$ (fig. 2C).

*6) SD_lograte_0.55*

Identical to the focal scheme, but with the variance of the logarithm of rate equal to $0.55^2$ instead of $0.2^2$, such that $\frac{\text{sd(rate)}}{\text{mean(rate)}} \approx \frac{0.015}{0.025} = 0.6$ (fig. S9C).

#### Sequence length

*7) 100-aa*

Identical to the focal scheme, but with a simulated alignment length of 100 amino acids (figs. S7 and S8).

*8) 500-aa*

Identical to the focal scheme, but with a simulated alignment length of 500 amino acids (figs. S7 and S8).

*9) 1000-aa*

Identical to the focal scheme, but with a simulated alignment length of 1000 amino acids (figs. S7 and S8).

*10) 2000-aa*

Identical to the focal scheme, but with a simulated alignment length of 2000 amino acids (figs. S7 and S8).

#### Tip number

*11) 40-tip*

Identical to the focal scheme, but with a simulated phylogeny of 40 tips (fig. S10A).

*12) 60-tip*

Identical to the focal scheme, but with a simulated phylogeny of 60 tips (fig. S10B).

*13) 80-tip*

Identical to the focal scheme, but with a simulated phylogeny of 80 tips (fig. S10C).

*14) 100-tip*

Identical to the focal scheme, but with a simulated phylogeny of 100 tips (fig. S10D).

#### Substitution model

In the following schemes, everything was the same as used in the focal simulation scheme except for different substitution models.

*15) LG+C60+G{0.5} (Gamma_0.5)*

LG substitution model plus C60 amino acid site frequency profiles and among-site relative rate variation under a four-category discrete gamma distribution with $\alpha = 0.5$ (fig. 2A).

*16) LG+C60+G{1.5} (Gamma_1.5)*

LG substitution model plus C60 amino acid site frequency profiles and among-site relative rate variation under a four-category discrete gamma distribution with $\alpha = 1.5$ (fig. S9A).

*17) LG+G*

LG substitution model plus C60 amino acid site frequency profiles and among-site relative rate variation under a four-category discrete gamma distribution with $\alpha = 1$ (figs. 1-3).

*18) LG+G+I*

The model LG+G supplemented by a proportion of invariant sites (p_inv), which was drawn from Uniform$(0,1)$ (fig. 3C).

*19) LG+C60+G+I*

The model LG+C60+G+I supplemented by a proportion of invariant sites (p_inv), which was drawn from Uniform$(0,1)$ (fig. 3C).

*20) LG+R*

Free rate model (Soubrier et al., 2012; Yang, 1995) with eight rate categories, where category weights $w = (w_1, \dots, w_8)$ and unscaled positive base rates $r' = (r'_1, \dots, r'_8)$ were drawn from two independent, flat Dirichlet$(1, \dots, 1)$ distributions. The final, normalized relative rate for each category $i$, denoted by $r_i$, was then calculated as $r_i = \frac{r'_i}{(\sum_{j=1}^{8} w_j r'_j)}$. This guarantees $r_i > 0$ and $\sum_{i=1}^{8} w_i \times r_i = 1$ (fig. 3C).

*21) EX2+G*

Mixture substitution model EX2 (Le et al., 2008) consisting of two matrices corresponding to exposed/buried sites, plus among-site relative rate variation under a four-category discrete gamma distribution with $\alpha = 1.0$ (fig. S5A).

*22) EX3+G*

Mixture substitution model EX3 consisting of three matrices corresponding to highly exposed/intermediate/buri[illegible] sites Le et al. (2008), plus among-site relative rate variation under a four-category discrete gamma distribution with $\alpha = 1.0$ (fig. S5B).

*23) EX2+C20+G*

Mixture substitution model EX2 plus C20 amino acid site frequency profiles and among-site relative rate variation under a four-category discrete gamma distribution with $\alpha = 1.0$ (fig. S4A).

*24) EX3+C20+G*

Mixture substitution model EX3 plus C20 amino acid site frequency profiles and among-site relative rate variation under a four-category discrete gamma distribution with $\alpha = 1.0$ (fig. S4B)

*25) LG4M*

Substitution model LG4M (Le et al., 2012) and among-site relative rate variation under a four-category discrete gamma distribution with $\alpha = 1.0$ (fig. S5C).

### Note S2. Calibration information

#### S2.1 Simulation dataset

See "Calibrations used in molecular dating analysis on simulated datasets" in Materials and Methods.

#### S2.2 Simulation dataset with calibration misspecification

We also explored the impact of mis-specified calibrations on molecular-clock dating by shifting the calibrations backward or forward relative to their true ages (Figs. 3A–3B).

*1) $f0.2$, $f0.5$: forward*

Building upon the *root_only*, *single_interval*, and *two_intervals* calibration schemes (see Fig. 1 and Materials and Methods), we shifted the time bounds of all calibrations forward (**toward the present**) by proportions of 0.2 and 0.5 of their true values.

$$\begin{aligned} \text{Original:} \quad & T_{\text{calibrated\_node}} \sim \text{Uniform}\big((1-k)\,\text{true\_age}, (1+k)\,\text{true\_age}\big) \\ \text{Shifted:} \quad & T_{\text{calibrated\_node}} \sim \text{Uniform}\big((1-2k)\,\text{true\_age}, \text{true\_age}\big), \qquad k \in \{0.2, 0.5\}. \end{aligned}$$

*Example (f0.2).* for a calibration with true age 10 and original distribution $\text{Uniform}(8,12)$, applying the 20% forward shift yields: $\text{Uniform}(6,10)$. Thus, every calibration is truncated toward the present while maintaining the original upper bound at the true age.

*2) $b0.2$, $b0.5$: backward*

Building upon the *root_only*, *single_interval*, and *two_intervals* calibration schemes, we shifted every calibration window backward (**toward the past**) by proportions $k \in \{0.2, 0.5\}$ of the corresponding true node ages.

$$\begin{aligned} \text{Original:} \quad & T_{\text{calibrated\_node}} \sim \text{Uniform}\big((1-k)\,\text{true\_age}, (1+k)\,\text{true\_age}\big) \\ \text{Shifted:} \quad & T_{\text{calibrated\_node}} \sim \text{Uniform}\big(\text{true\_age}, (1+2k)\,\text{true\_age}\big), \qquad k \in \{0.2, 0.5\}. \end{aligned}$$

*Example ($b0.2$).* A calibration with true age 10 and original distribution $\text{Uniform}(8,12)$ becomes $\text{Uniform}(10,14)$ after the 20% backward shift.

#### S2.3 Empirical dataset

Microsporidia

Because we are aware of any fossil records within neither Microsporidia nor its closely related lineages, we set a single calibration at the root, which is the LCA of Microsporidia and Rozella (outgroup) by a uniform distribution $\text{Uniform}(0,0.9)$ (time unit: Ga). This is based on previous estimates, based on indirect evidence and without using a molecular clock approach, that Rozella split from Microsporidia 700 Ma (Chang et al., 2022) but likely within 900 Ma (0.9 Ga) (Chang et al., 2015). Alternative calibrations can be found in Data S6.

Rickettsiales

We applied the mitochondrial endosymbiosis strategy with eukaryotic fossils to date the evolution of Rickettsiales, the earliest-splitting order within Alphaproteobacteria. This is based on the mitochondrial endosymbiosis stating that mitochondria originated from a lineage closely related to Alphaproteobacteria. Specifically, we applied the “mito-encoded” dataset used in our previous study (Wang and Luo, 2021). This dataset consists of 24 mitochondrial genome-encoded genes that are shared with $\alpha$-Proteobacteria (Wang and Wu, 2015), as well as four within-eukaryote calibrations: total-group eudicots (250-125 Ma), total-group bryophytes (509-450 Ma), total-group Florideophyceae (root_max to 550 Ma), total-group Rhodophyta (root_max to 1047 Ma), all uniformly distributed. Note that the last calibration was originally placed as the crown-group Rhodophyta in (Wang and Luo, 2021), but we changed it to total-group Rhodophyta in the present study based on recent arguments about the most reasonable placement of the fossils (Betts et al., 2018; Gibson et al., 2018). In addition, we applied another calibration within the outgroup: total-group Chromatiaceae (root_max to 1640 Ma) based on a biomarker interpreted as evidence for Chromatiaceae in several genera of phototrophic purple sulfur bacteria (Chromatiaceae, $\gamma$-Proteobacteria) (Brocks and Schaeffer, 2008), as used in several studies (Battistuzzi and Hedges, 2009; Wang and Luo, 2025). The maximum of the age of the root (the LCA of $\alpha-, \beta-, \gamma-$Proteobacteria) was arbitrarily set to 3000 Ma, which is conservative enough based on prior studies (Wang and Luo, 2025; **?**). Alternative calibrations can be found in Data S6.

### Note S3. Supplemental methods

#### S3.1 Substitution model selection for the empirical dataset

We selected the best-fitting substitution models as the one that achieved the lowest AIC (Akaike Information Criterion) value using ModelFinder (Kalyaanamoorthy et al., 2017) implemented in IQ-Tree. The commands were given as follows.

```
iqtree -s 0/combined.aln -m TESTONLY -mset
 ↪  LG,LG4M,LG4X,EHO,EX2,EX3,EX_EHO,UL2,UL3 -mrate
 ↪  E,G,I,G+I,R1,I+R1,R4,I+R4 -T 1 -redo -pre iqtree -quiet
```

```
iqtree -s combined.aln -m TESTONLY -mset
 ↪  LG+C10,LG+C20,LG+C30,LG+C40,LG+C50,LG+C60,C10,C20,C30,C40,C50,C60
 ↪  -mrate E,G,I,G+I,I+R1,R4,I+R4 -T 10 -redo -pre iqtree_mwopt -mwopt
 ↪  -quiet
```

Note that “+R1” in the substitution model LG+Cxx+R1 is to turn off the +G automatically implied by Cxx by forcing the number of the relative rate categories in the so-called free rate model (+R) to be 1.0 (in this way, +R1 will “overwrite” +G4).

#### S3.2 MCMCtree analysis

For empirical data analysis, we set the parameters in MCMCtree molecular clock analysis following our previous study (Wang and Luo, 2025). The time unit was 100 Myr (0.1 Gyr) . The divergence time priors for uncalibrated nodes were based on a birth-death process (Nee et al., 1994; Yang and Rannala, 1997) with parameters: birth rate = 1, death rate = 1, and taxon sampling proportion = 0.001, resulting in a (nearly) uniform time prior (Dos Reis et al., 2015). Rate variation across branches was modeled with a gamma distribution, G(1,10). The rate prior was a diffuse Dirichlet-gamma prior centered at 0.02 substitutions per site per 0.1 Gyr with the shape value $\alpha = 1$ and a scale parameter $\beta = 50$. Each MCMC chain was run, sampling every 500 iterations to get 1000 posterior samples, following a burn-in period of 50000 iterations. The above guarantees the convergence as indicated by effective sample size (ESS) larger than 200.

For the simulation analysis, the parameters of the MCMCtree analysis were set according to the true parameters used in simulation. Each MCMC chain was run, sampling every 10 iterations to get 500 posterior samples, following a burn-in period of 1000 iterations, which is enough to guarantee convergence with ESS $\geq$ 100 given the small dataset (20 tips and 300 amino acids).

#### S3.3 More details on the 4-th order finite difference method to calculate the Hessian

The multivariate Taylor expansion of $\ell(\theta)$ around the point $(\hat{\theta}_i, \hat{\theta}_j)$ is

$$\ell\big(\hat{\theta}+e_i a+e_j b\big) = \ell + \sum_{k=1}^{5} \frac{1}{k!} \sum_{m=0}^{k} \binom{k}{m} a^m b^{k-m} \frac{\partial^k \ell}{\partial \theta_i^m \partial \theta_j^{k-m}} + \mathscr{O}\big(a^6, b^6\big).$$

Evaluating $\ell(\hat{\theta})$ at $1+4\times 4 = 17$ symmetric points

$$(\hat{\theta}_i, \hat{\theta}_j), \quad (\hat{\theta}_i \pm h, \hat{\theta}_j \pm h), \quad (\hat{\theta}_i \pm 2h, \hat{\theta}_j \pm 2h),$$

and their combinations, while holding all other variables $\hat{\theta}_k$ for $k \neq i, j$ constant, we construct a linear system $Ax = B$. In this system, $A \in \mathbb{R}^{21\times 17}$ is the matrix of Taylor coefficients for derivatives up to 5th order, $x \in \mathbb{R}^{17}$ is the vector of weights for the function evaluations, and $B \in \mathbb{R}^{17}$ is the target vector encoding the desired derivative and cancellation of higher-order terms (Note S3). The row size of $A$ is 21 because there are 21 items in the expansion (Term 1) to eliminate to achieve an error order of $\mathscr{O}(h^4)$.

Specifically, the Taylor coefficient matrix $\mathbf{A} \in \mathbb{R}^{21\times 17}$ is constructed as:

$$\mathbf{A} = \begin{bmatrix} 1 & 1 & 1 & \cdots & 1 \\ 0 & k_1 & k_2 & \cdots & k_{17} \\ 0 & l_1 & l_2 & \cdots & l_{17} \\ 0 & \frac{k_1^2}{2} & \frac{k_2^2}{2} & \cdots & \frac{k_{17}^2}{2} \\ 0 & k_1 l_1 & k_2 l_2 & \cdots & k_{17} l_{17} \\ \vdots & \vdots & \vdots & \ddots & \vdots \\ 0 & \frac{l_1^5}{120} & \frac{l_2^5}{120} & \cdots & \frac{l_{17}^5}{120} \end{bmatrix}_{21\times 17}$$

where $(k_i, l_i)$ are the 17 evaluation point coordinates:

$$\begin{aligned} &\{(0,0)\} \quad \text{(center point)} \\ &\cup\{(\pm 1, \pm 1)\} \quad \text{(diagonal at } h\text{)} \\ &\cup\{(\pm 2, \pm 2)\} \quad \text{(diagonal at } 2h\text{)} \\ &\cup\{(\pm 1, \pm 2)\} \quad \text{(mixed: } h \text{ in } x\text{, } 2h \text{ in } y\text{)} \\ &\cup\{(\pm 2, \pm 1)\} \quad \text{(mixed: } 2h \text{ in } x\text{, } h \text{ in } y\text{)} \end{aligned}$$

The target vector $\mathbf{B} \in \mathbb{R}^{17\times 1}$ is given as

$$\begin{aligned} \mathbf{B}_{1\times 17}^{\top} &= \begin{bmatrix} 0 & 0 & 0 & 0 & 1 & 0 & \cdots & 0 \end{bmatrix} \quad (\text{for } H_{ij}), \\ \mathbf{B}_{1\times 17}^{\top} &= \begin{bmatrix} 0 & 0 & 0 & 1 & 0 & \cdots & 0 \end{bmatrix} \quad (\text{for } H_{ii}). \end{aligned}$$

The system is underdetermined, as its rank is 15, smaller than the number of equations (which is 17). This leads to infinitely many solutions. To find the solution that minimizes $\|x\|_F^2$, we use

$$x = A^{\mathsf{T}}\left(AA^{\mathsf{T}}\right)^{-1} B,$$

where $A^{\mathsf{T}}\left(AA^{\mathsf{T}}\right)^{-1}$ is the right pseudoinverse of $A$.

Denote

$$\ell_{k,j} = \ell\big(\hat{\theta} + khe_i + jhe_j\big).$$

These give the approximations

$$H_{ii}(\hat{\theta}) \approx \frac{1}{h^2}\left(-\tfrac{5}{4}\Delta_{0,0} + \tfrac{1}{3}\Delta_{1,1} - \tfrac{1}{48}\Delta_{2,2} + \tfrac{12}{169}\Delta_{1,2} - \tfrac{12}{169}\Delta_{2,1}\right), \tag{1}$$

$$H_{ij}(\hat{\theta}) \approx \frac{1}{600h^2}\left(74\Delta_{1,1} + 44\Delta_{2,2} - 63\Delta_{1,2} + 63\Delta_{2,1}\right), \tag{2}$$

where

$$\begin{cases} \Delta_{1,1} = \ell_{1,1} - \ell_{1,-1} - \ell_{-1,1} + \ell_{-1,-1}, \\ \Delta_{2,2} = -\ell_{2,2} + \ell_{2,-2} + \ell_{-2,2} - \ell_{-2,-2}, \\ \Delta_{1,2} = \ell_{1,2} - \ell_{1,-2} - \ell_{-1,2} + \ell_{-1,-2}, \\ \Delta_{2,1} = \ell_{2,1} - \ell_{2,-1} - \ell_{-2,1} + \ell_{-2,-1}. \end{cases}$$

R Code:

```
build_A_matrix <- function() {
    steps <- data.frame(
        k = c(1, 1, -1, -1,
              2, 2, -2, -2,
              1, 1, -1, -1,
              2, 2, -2, -2),
        l = c(1, -1, 1, -1,
              2, -2, 2, -2,
              2, -2, 2, -2,
              1, -1, 1, -1)
    )

    # for (0,0)
    steps <- rbind(data.frame(k = 0, l = 0), steps)

    if (nrow(steps) != 17) stop("Incorrect number of evaluation points")

    A <- matrix(0, nrow = 21, ncol = 17)

    for (i in 1:17) {
        k <- steps$k[i]
        l <- steps$l[i]

        A[, i] <- c(
            1,
            k, l,
            k^2/2, k*l, l^2/2, # (f_xx, f_xy, f_yy)
            k^3/6, k^2*l/2, k*l^2/2, l^3/6, # 3rd
            k^4/24, k^3*l/6, k^2*l^2/4, k*l^3/6, l^4/24, # 4th
            k^5/120, k^4*l/24, k^3*l^2/12, k^2*l^3/12, k*l^4/24, l^5/120
             ↪  # 5th
        )
    }

    return(A)
}


compute_pseudo_inverse_solution <- function(A, B) {
  AAT <- A %*% t(A) + 1e-10 * diag(nrow(A))
```

```
  # Right pseudo-inverse: A^T (A A^T)^{-1}
  A_pinv <- t(A) %*% solve(AAT)
  x <- A_pinv %*% B
  return(x)
}


build_B_vector <- function(placement="ij") {
    B <- matrix(0, nrow = 21, ncol = 1)
    if(placement == 'ij'){
        B[5, 1] <- 1
    }
    if(placement == 'ii'){
        B[4,1] <- 1 # B[6,1] gives the same result as it's symmetric
    }
    return(B)
}


##################################################################
A <- build_A_matrix()
# H_ij
B <- build_B_vector("ij"); compute_pseudo_inverse_solution(A,B)
# H_ii
B <- build_B_vector("ii"); compute_pseudo_inverse_solution(A,B)
```
